# An insight into neurotoxic and toxicity of spike fragments SARS-CoV-2 by exposure environment: A threat to aquatic health?

**DOI:** 10.1101/2021.01.11.425914

**Authors:** Ives Charlie-Silva, Amanda P. C. Araújo, Abraão T. B. Guimarães, Flávio P Veras, Helyson L. B. Braz, Letícia G. de Pontes, Roberta J. B. Jorge, Marco A. A. Belo, Bianca H V. Fernandes, Rafael H. Nóbrega, Giovane Galdino, Antônio Condino-Neto, Jorge Galindo-Villegas, Glaucia M. Machado-Santelli, Paulo R. S. Sanches, Rafael M. Rezende, Eduardo M. Cilli, Guilherme Malafaia

## Abstract

The Spike protein (S protein) is a critical component in the infection of the new coronavirus (SARS-CoV-2). The objective of this work was to evaluate whether peptides from S protein could cause negative impact in the aquatic animals. The aquatic toxicity of SARS-CoV-2 spike protein peptides derivatives has been evaluated in tadpoles (n = 50 tadpoles / 5 replicates of 10 animals) from species Physalaemus cuvieri (Leptodactylidae). After synthesis, purification, and characterization of peptides (PSDP2001, PSDP2002, PSDP2003) an aquatic contamination has been simulatedwith these peptides during 24 hours of exposure in two concentrations (100 and 500 ng/mL). The control group (“C”) was composed of tadpoles kept in polyethylene containers containing de-chlorinated water. Oxidative stress, antioxidant biomarkers and neurotoxicity activity were assessed. In both concentrations, PSPD2002 and PSPD2003 increased catalase and superoxide dismutase antioxidants enzymes activities, as well as oxidative stress (nitrite levels, hydrogen peroxide and reactive oxygen species). All three peptides also increased acetylcholinesterase activity in the highest concentration. These peptides showed molecular interactions in silico with acetylcholinesterase and antioxidant enzymes. Aquatic particle contamination of SARS-CoV-2 has neurotoxics effects in P. cuvieri tadpoles. These findings indicate that the COVID-19 can constitute environmental impact or biological damage potential.

**HIGHLIGHTS:**

- SARS-CoV-2 spike protein peptides (PSDP) were synthesized, purified, and characterized by solid phase peptide synthesis.
- PSDP peptides promoted REDOX imbalance and acute neurotoxicity in tadpoles (Physalaemus cuvieri)
- In silico studies have shown interactionsbetween peptides and acetylcholinesterase and antioxidant enzymes
- Aquatic particle contamination of SARS-CoV-2 can constitute additional environmental damage

**GRAPHICAL ABSTRACT:** 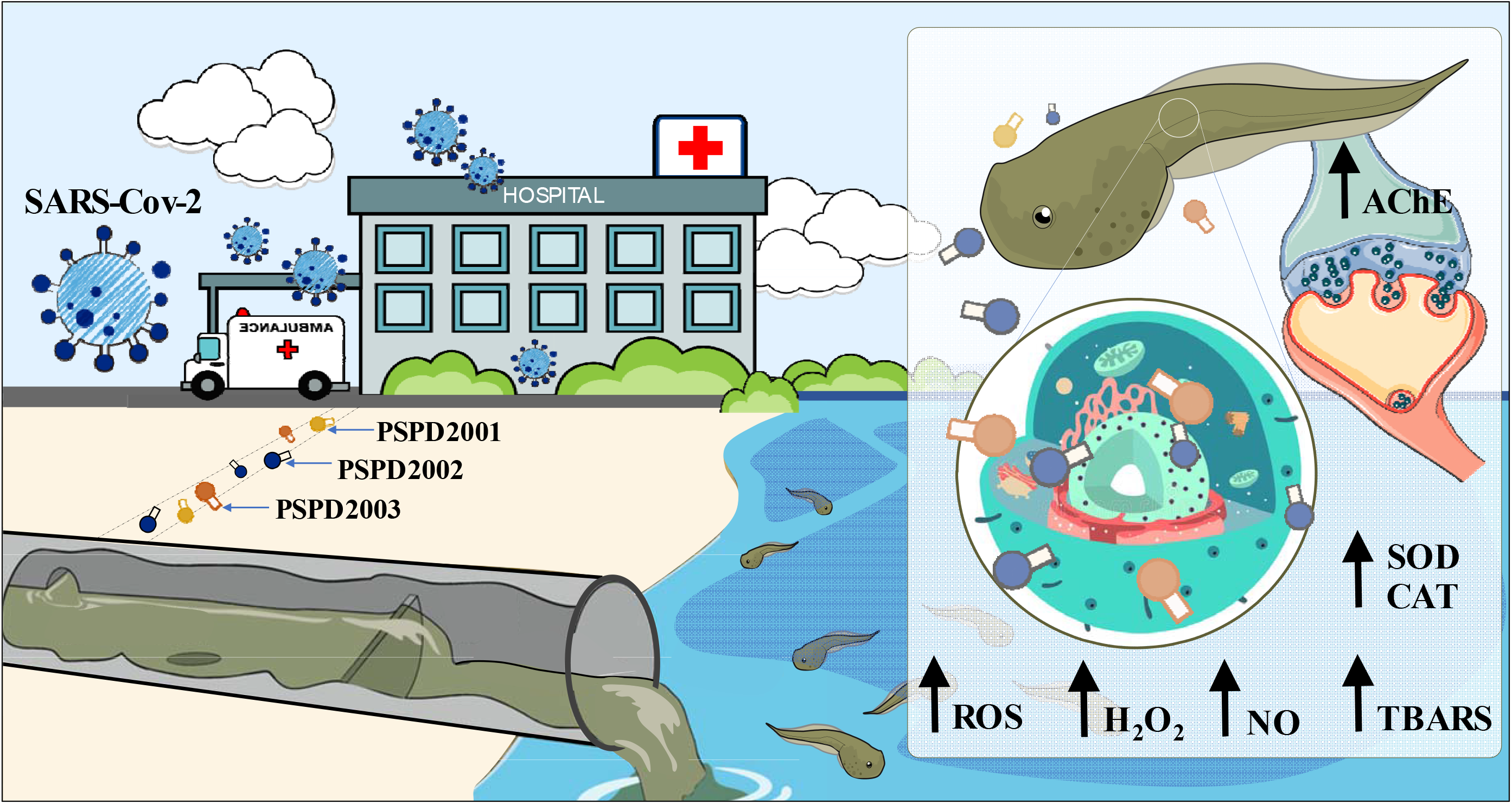

## 1. INTRODUCTION

Coronavirus Disease-2019 (COVID-19) pandemic, caused by SARS-CoV-2 (Severe acute respiratory syndrome coronavirus 2), an emergent beta-coronavirus threatening human health, has led to a dramatic worldwide crisis and presents unprecedented global challenges on everybody’s daily life, social aspects, political affairs, and health measures (Chakraborty & Prasenjit, 2020)

Remarkably, the poor and the most vulnerable people are at critical high risk, and Oxfam, an international confederation of 20 NGO’s already calculates that nearly 500 million people worldwide may succumb to poverty resulting from the same (OI, 2020). By Dec. 27, over 79.2 million cases and over 1.7 million deaths have been reported since the start of the pandemic (FAO, 2020). Resulting from the same, in only 12 months, we have learned a lot about SARS-CoV-2. Our ability to test for and manage COVID-19 has improved, but ongoing debate remains about how SARS-CoV-2 is transmitted (Editorial Lancet, 2020).

The most recurrent forms of SARS-CoV-2 transmission are through direct contact with an infected person (Meyerowitz et al., 2020), inhaling respiratory droplets containing the virus (Harrison et al., 2020), or accessing a contaminated environment where suspended particles are present over longer distances and time than droplet transmission (Graham et al. 2020).

However, by reviewing the environmental threats of the virus reported so far, it is concluded that the virus can survive on inanimate surfaces such as metal, glass, or plastic for up to 9 days if any effective disinfection procedure with ozone, ethanol, hydrogen peroxide, UV light, chlorine or its derivatives as sodium hypochlorite is not conducted in between (Kampf et al. 2020).

Although the direct contact described above concerns meaningful, a different environmental transmission source with the virus now recognized is the indirect contact through the infected people’s stool and urine (Chen et al., 2020; Xiao et al., 2020; Jones et al., 2020). Unconventional studies support this notion by reporting positive SARS-CoV-2 viral titers in domestic sewages (Pandey et al., 2020; Elsamadony et al., 2020; Polo et al., 2020). So far, Sars-Cov-2 has been detected in several countries wastewaters of the American, European, and Asian continents, suggesting as mandatory the monitoring of the secondary transmission of the new coronavirus via wastewater (Liu et al. 2020). On a more compelling perspective, strong evidence suggests that surveillance of primarily settled solids in wastewater through one-step ddPCR is a solid strategy to track the spread of Covid-19 disease transmission before the clinical cases break out in a particular location (Graham, 2020).

Moreover, following this trend can shed light on the characteristics of infection that are difficult to capture in clinical investigations, such as the dynamics of infection and early viral elimination (Wu et al., 2020).

The increase in the generation of household waste (Sharma et al., 2020; Zand & Heir, 2020; Urban et al., 2021), hospitals (Abu-Qdais et al., 2020; Sangkham, 2020; Yang et al., 2020), and notable civil buildings (Carvalho et al., 2020; Abu-Rayash et al., 2020; Santiago et al., 2020) constitute some of the environmental grounds where no information on the ecotoxicological effects of SARS-CoV-2 proteic or genetic structural components impact on freshwater vertebrates exists.

Therefore, this lack of knowledge requires urgent attention by developing studies to assess how COVID-19 impacts the aquatic populations in the close vicinity of the anthropogenic activities described above. Such studies may focus on supporting actions or strategies on the remediation or at least mitigation of impacts in favor of conserving nontarget species at the edge of any Sars-Cov-2 variant.

The Spike (S) protein is a critical component of the new Sars-Cov-2 coronavirus found on the surface of the SARS-Cov-2 virus, giving it a “crown” appearance. The S protein is a granule-shaped structural protein with a length of about 1200 aa, which helps the virus bind to cell receptors and mediates viral infection and pathogenesis (Coughlan, 2020). The S protein plays a key role in the receptor recognition and cell membrane fusion process with ACE-2 (angiotensin-converting enzyme 2) (Huang et al., 2020). Therefore, it is not surprising this ligand-receptor interaction of the S protein is the primary target to produce vaccines against COVID-19, as reported in different studies (Bangaru et al., 2020; Samrat et al., 2020; Keech et al., 2020; Yang et al., 2020; Qi et al., 2020; Ravichandran et al., 2020).

Several in vivo platforms to dissect the cellular and molecular programs governing Sars-Cov-2 viral dissemination on vertebrates are available. However, the number of aquatic model animals that may support trials and provide reliable information is almost inexistent. Among them, the zebrafish model represents an attractive model to explore the desired effects on the context of a full vertebrate (Galindo-Villega, 2020). Unfortunately, the zebrafish has not been vigorously infected in vivo trials so far by the causative agent of Covid-19 (Gaudin & Goetz, 2021).

Following a synthetic approach in previous research, we have developed three peptides of the full-length SARS-CoV-2 Spike protein (PSPD2001, PSPD2002, and PSPD 2003) after a pattern memorization phagolysosomal proteolysis (Fernandes et al. 2020). To attempt to elucidate whether and how the Sars-Cov-2 influence the aquatic animals, in this study, we investigate the same by adding the three produced synthetic peptides to mimic the resulting Covid-19 aquatic contamination in wastewater. The tadpole *Physalaemus cuvieri*, is a prevalent amphibian species found in many freshwater habitats throughout Brazil and South America ((Miranda et al., 2019; Herek et al., 2020; Araújo et al., 2020ab; Rutkoski et al., 2020). Its population stability and abundance in the areas that occur (Frost 2017), good adaptability in the laboratory, and early biological response justify the species’ choice (Herek et al., 2020; Araújo et al., 2020ab; Rutkoski et al., 2020). Previous studies using this species report the effects of water pollution caused by wastewater runoff (Wrubleswski et al. 2018). Therefore, in this study, we selected *P. cuvieri* as our choice of a translational model vertebrate.

From different biomarkers indicative of an imbalance in oxidation-reduction (REDOX) and neurotoxicity processes, we aimed to test the hypothesis that nanometric concentrations of the SARS-CoV-2 Spike protein fragments in water may affect the health of amphibians. We believe that studies like ours are needed not only to expand our knowledge about the impacts of COVID-19 on aquatic biodiversity; but also to predict the environmental impacts of the recent pandemic on the populations of neotropical amphibians, which have already, over the years, shown a drastic population decline (Pechmann et al., 1991; Blaustein et al., 2002; Ranvestel et al., 2004; Grant et al., 2020).

## 2. MATERIAL AND METHODS

### 2.1. Synthesis, purification, and characterization of peptides

#### 2.1.1. Synthesis of SARS-CoV-2 Spike protein peptides

The peptides were obtained manually using the solid phase peptide synthesis method (SPFS) using the Fmoc strategy (Raibaut et al., 2014; Behrendt et al., 2016). The couplings were carried out by activating the carboxyl groups of the Fmoc-amino acids with a solution of diisopropylcarbodimide and hydroxybenzotriazole (HOBT), for2 h. In this step, a 2-fold excess of Fmoc-amino acids and coupling agents in relation to the number of reactive sites in the resin was used. Deprotection of the amino group after coupling, i.e., removal of the base labile Fmoc group was carried out by reaction with a 20% solution of 4-methyl-piperidine in dimethylformamide (DMF) following the exit of the protective group through the colorimetric test ninhydrin (Luna et al., 2016), which identifies free amine groups converting the yellow solution to violet-blue after incubation at 110°C for 3 min. The resins used for synthesis were Fmoc-Cys (Trt)-Wang, Fmoc-Thr (TBu)-Wang, and Fmoc-Asn (Trt)-Wang for peptides Arg-Val-Tyr-Ser-Ser-Ala-Asn-Asn-Cys-COOH (PSPD2001); Gln-Cys-Val-Asn-Leu-Thr-Thr-Arg-Thr-COOH (PSPD2002) and Asn-Asn-Ala-Thr-Asn-COOH (PSPD2003), respectively

#### 2.1.2. Cleavage of SARS-CoV-2 Spike protein peptides

After coupling all the amino acid residues from the peptide sequences, the chains were removed from the solid support by acid cleavage using trifluoroacetic acid (TFA) for 2 h [similarly to Guy & Fields (1997)]. In addition to TFA, reaction suppressors were added according to the sequence of each peptide. After cleavage, the peptides were precipitated with cold ether and later extracted with 0.045% TFA solution in purified water. The solutions were lyophilized to obtain solid crude material.

#### 2.1.3. Purification of SARS-CoV-2 Spike protein peptides

The crude compounds were purified by high-performance liquid chromatography (HPLC) with a reverse-phase column using different purification methods according to the retention time obtained in a gradient program of 5 to 95% in 30 min (exploration gradient) in Analytical HPLC (Klaassen et al.,2019). Table S1 (see “Supplementary Material”) presents a summary of the purification methods adopted in our study. The purification solvents were water containing 0.045% TFA (solvent A) and acetonitrile containing 0.036% TFA (solvent B).

After collecting, lyophilizing, and weighing the pure material fractions, its yield was calculated, obtaining 13.0% for PSPD2001, 21.4% for PSPD2002, and 18.2% for PSPD2003. The pure and solid material was subjected to chromatographic analysis to determine the purity of the final product. Only compounds with purity equal to or greater than 95% were considered for biological analysis,

#### 2.1.4. Characterization of SARS-CoV-2 Spike protein peptides

The analysis of the synthesized peptides’ identity was carried out in a mass spectrometer (Metzgeret al., 1994)Thermo LCQ-fleet, with ESI-IT-MS configuration. For this, the sample solutions were directly infused at a concentration of approximately 10 mg/L in acetonitrile/water containing 0.1% v/v formic acid. The infusion rate was adjusted to 5.0 μL/min, and the electrospray source was operated in a positive mode, applying 4.5 kV to the electrospray capillary.

### 2.2. Alignment of SARS-CoV-2 Spike protein peptides

The similarities between the PSPD2001, PSPD2002 and PSPD2003 peptides synthesized in the present study were tested using the CLUSTAL W version 1.83 software [Higgins et al. (1996), Pais et al. (2014) – http://www.ebi.ac.uk/clustalw/]. The peptides were aligned with proteins deposited in the NCBI/BLAST (Basic Local Alignment Search Tool), consisting of a set of programs that look for similarities between different sequences. The investigated sequences’ alignment was carried out with the nucleic acid and/or protein database (http://www.ncbi.nlm.nih.gov/blast). Within BLAST, the search was carried out in the “Protein blast” using as a database the “Swissprot protein sequence (swissprot)”, algorithm – blastp (protein BLAST) and the search was restricted to Physalaemus cuvieri (taxid:218685). The database UniProtKBSwissProt (http://www.uniprot.org/) was used to obtain detailed information on the protein aligning with the selected peptides revealed.

### 2.3. Model system and experimental design

To evaluate the synthesized peptides’ aquatic toxicity, we used tadpoles of the species Physalaemus cuvieri (Leptodactylidae) as a model system. All tadpoles used came from an ovigerous mass (containing approximately 1500 eggs), according to Pupin et al. (2010). They were collected in a lentic environment (Urutaí, GO, Brazil) surrounded by native vegetation from the Cerrado biome, under license no. 73339-1 of the Biodiversity Authorization and Information System (SISBIO/MMA/ICMBio) in Brazil.

Upon arrival at the laboratory, the eggs were kept in an aquarium (40.1 x 45.3 x 63.5 cm) containing 80 L of naturally dechlorinated water (for at least 24 h), under controlled light conditions (light-dark cycle, 12:12 h), temperature (26 °C ± 1 °C – similar to the natural environment) and constant aeration (maintained by air compressors), being fed once a day (ad libitum) with commercial fish food (guarantee levels: 45% crude protein, 14% ether extract, 5% crude fiber, 14% mineral matter and 87% dry matter). After the eggs hatched, the tadpoles remained in these conditions until they reached stage 27G, according to Gosner (1960) (body biomass: 70 mg ± 4.1 mg and total length: 20.1 mm ± 0.7 mm – mean ± SEM). The healthy tadpoles (i.e., with normal swimming movements and without morphological deformities or apparent lesions) were divided into seven experimental groups (n = 50 tadpoles / each – 5 replicates composed of 10 animals/each. The control group (“C”) was composed of tadpoles kept in polyethylene containers containing 50 ml of de-chlorinated water, free of any peptide. The animals kept in water containing the peptides comprised the groups “PSPD2001”, “PSPD2002”, and “PSPD2003”. Two concentrations were tested for each peptide (100 and 500 ng/mL, defined based on reports that SARS-CoV-2 in sweet environments occurs in minimal concentrations (Shutler et al., 2020; Guerrero-Latorre et al., 2020; Tran et al., 2020). The exposure period (24 h; in the static system) was defined considering the low persistence of SARS-CoV-2 in the aquatic environment after being released with human feces (Bivins et al., 2020).We emphasize that, throughout the exposure period, different physical-chemical parameters of the quality of the exposure waters were monitored (every 6 hours), keeping them equitable between treatments (temperature: 23°C ± 1.14; atmospheric pressure (atm): 0.91 ± 0.0001; electrical resistivity (Ωm): 0.01 ± 0.0001; electrical conductivity (μS/cm^2^): 96.2 ± 1.83; total dissolved solids (mg/L): 48, 2 ± 0.83; salinity: 0.04 ± 0.004; oxidation-reduction potential (ORP): 130.21 ± 6.17; dissolved oxygen (mg/L): 7.72 ± 0.78 and pH: 7, 2 ± 0.38).

### 2.4. Toxicity biomarkers

We evaluated peptide-induced toxicity from predictive biomarkers of REDOX imbalance and neurotoxicity after exposure, considered classic and essential parameters in ecotoxicological studies (Valavanidis et al., 2016). For this, pools of four animals/each composed the samples to be analyzed. Such animals were weighed and later macerated in 1 mL of phosphate-buffered saline (PBS), centrifuged at 13.000 rpm for 5 min (at 4°C). Thesupernatant was separated into aliquots to be used in different biochemical evaluations. Entire bodies were used in the experiment due to the hard time isolating specific organs from small animals. Organ-specific biochemical assessment in tadpole requires highly accurate dissection due to their small size, making it difficult to process large sample numbers under a time constraint (Khan et al. 2015). Organ “contamination” by organic matter and/or by other particles consumed by tadpole can be biased at biochemical analysis applied to organs at dissection time (Lusher et al. 2017; Guimarães et al., 2021).

#### 2.4.1. REDOX state

##### 2.4.1.1. Oxidative stress biomarkers

The effects of exposure to peptides on oxidative stress reactions were evaluated based on indirect nitric oxide (NO) determination on REDOX regulated processes via nitrite measurement (Soneja et al. 2005); thiobarbituric acid reactive species (TBARS), a predictive of lipid peroxidation (De-Leon & Borges, 2020); production of reactive oxygen species (ROS) and on hydrogen peroxide (H_2_O_2_), which plays an essential role in responses to oxidative stress in different cell types (Sies, 2017). The Griess colorimetric reaction was used to measure NO (Grisham et al., 1996). This reaction consisted of detecting nitrite resulting from NO oxidation. TBARS levels were determined based on procedures described by Ohkawa et al. (1979) and modified by Sachettet al. (2020), with adaptations for conduction in microtubes and ELISA microplate reading. The reagent of 1.1,3.3-tetra-ethoxy-propane was used as a standard solution in the reaction with thiobarbituric acid (TBA) reactive substance, according Ro et al. (2020). In brief, this method’s principle depends on the Determination of the pink color produced by TBA interaction with malondialdehyde (MDA). The production of hydrogen peroxide and ROS was evaluated using the methodologies proposed by Elnemma et al. (2004) and Maharajan et al. (2018).

##### 2.4.1.2. Antioxidant response biomarkers

The activation or suppression of antioxidant activity by peptides was assessed by determining catalase and superoxide dismutase (SOD) enzyme activities, which are considered critical first-line antioxidants for defense strategies against oxidative stress (Ighodaro&Akinloye, 2018; Jing et al., 2020). While catalase activity was assessed based on Sinha et al. (1972) andMontalvão et al. (2021). The SOD was determined according to the method described by Del-Maestro & McDonald (1985) (initially) and adapted by Estrela et al. (2021). To assess the balance between the synthesis of hydrogen peroxide by SOD and its decomposition by catalase, the SOD/CAT ratio was calculated and recorded, as Liu et al. (2017).

#### 2.4.2. Neurotoxicity

Peptide neurotoxicity was assessed by quantifying acetylcholinesterase (AChE) activity, based on the method by Ellman et al. (1961), with minor detailed modifications in Estrela et al. (2021). AChE activity is used as a cholinergic function marker since it regulates the acetylcholine (ACh) amount interacting with its receptors (Tougu, 2001).

#### 2.4.3. Determination of the protein level

All results of the biochemical analyzes were expressed by the “g of proteins” of the samples. In this case, the protein level was determined with a commercial kit (Bioténica Ind. Com. LTD, Varginha, MG, Brasil. CAS number: 10.009.00) based on biuret reaction (Gornall et al., 1949; Henry et al., 1957). In general, Cu^2+^ions, in an alkaline medium, react with the peptide bonds of proteins forming the blue complex specifically with protein, and the intensity of color, measured by an ELISA reader at a wavelength of 492 nm, is proportional to the protein concentration.

### 2.5. Bioinformatics *in silico* analysis

Seeking to predict the binding mode and affinity of the bonds between the peptides synthesized in our study and the protein structures of the enzymes AChE, catalase, and SOD, we performed docking and chemoinformatic screens (Kolb et al., 2009). For this, we obtained the peptide ligand PSPD2002 and PSPD2003 in three dimensions through the web server PEP-FOLD3 (https://bioserv.rpbs.univ-paris-diderot.fr/services/PEP-FOLD3/). Protein structures and sequences of the P.cuvieri (i.e.:Leptodactylidae) taxonomic family were not found in the biological structure databases. Therefore, we use as target structures those from the Xenopodinae family, a family phylogenetically close to the group of Leptodactylidae (Jetz & Pyron, 2018). The AChE and catalase enzymes’ structures were obtained using the homology construction technique with values of similarity 65.48% and 87.14% to structures (targets) used for comparative modeling on the server SWISS-MODEL (https://swissmodel.expasy.org/), respectively. On the other hand, the structure of the SOD was obtained by Research Collaboratory for Structural Bioinformatics protein databank (https://www.rcsb.org) PDB code: 1XSO with 1.49Å resolution, obtained by X-ray diffraction of Xenopodinae origin. For molecular docking simulations, AutoDock tools (ADT) v4.2 (Morris et al., 2009) and AutoDock Vina 1.1.2 (Trott & Olson, 2010) were used. The procedure was carried out by removing water molecules and other residues present in the target structures. A polar hydrogen group was added to establish hydrogen bonds between the macromolecule and the ligand tested. The grid box was chosen based on the native ligand of the macromolecules (targets). The binding potency (ΔG affinity) was used to determine better molecular interactions. The results were visualized using ADT and UCSF Chimera X (Pettersen et al., 2021).

### 2.6. Data analysis

GraphPad Prism Software Version 8.0 (San Diego, CA, USA) was used to perform the statistical analysis. Initially, data were checked for deviations from the normality of variance and homogeneity of variance before analysis. Normality of data was assessed using the Shapiro-Wilks test, and homogeneity of variance by the Bartlett’s test. Multiple comparisons were performed using a one-way ANOVA and Tukey’s posthoc analysis (for parametric data) or the Kruskal-Wallis test, with Dunn’s posthoc (for non-parametric data). Correlation analysis was performed through Pearson’s (parametric data) or Spearman’s method (non-parametric data). Besides, the regression analysis was performed when significant differences were detected between different treatments. Levels of significance were set at (p) less than 0.05, 0.01, or 0.001.

## 3. RESULTS AND DISCUSSION

### 3.1. Synthesis and characterization of SARS-CoV-2 Spike peptides

Our study’s first stage was to synthesize and characterize the SARS-CoV-2 Spike peptides arbitrary named PSPD2001, PSPD2002, and PSPD2003. During the peptides’ cleavage, we performed the addition of different reaction suppressors to avoid the return of the side chain protectors present in some amino acids with a reactive side chain. The results obtained in this step are shown in Table 1. Regarding the mass spectrometry analysis, the spectra obtained indicated the molecular mass/charge ratio (m/z) of the identified compounds, allowing us to confirm the deprecated molecules’ achievement. The spectra can be seen in Figure S1 (see “Supplementary Material”), and Table 2 summarizes the results obtained in this step. Figure 1 also shows the structural models of the PSPD2001, PSPD2002, and PSPD2003.

**Figure 1.**
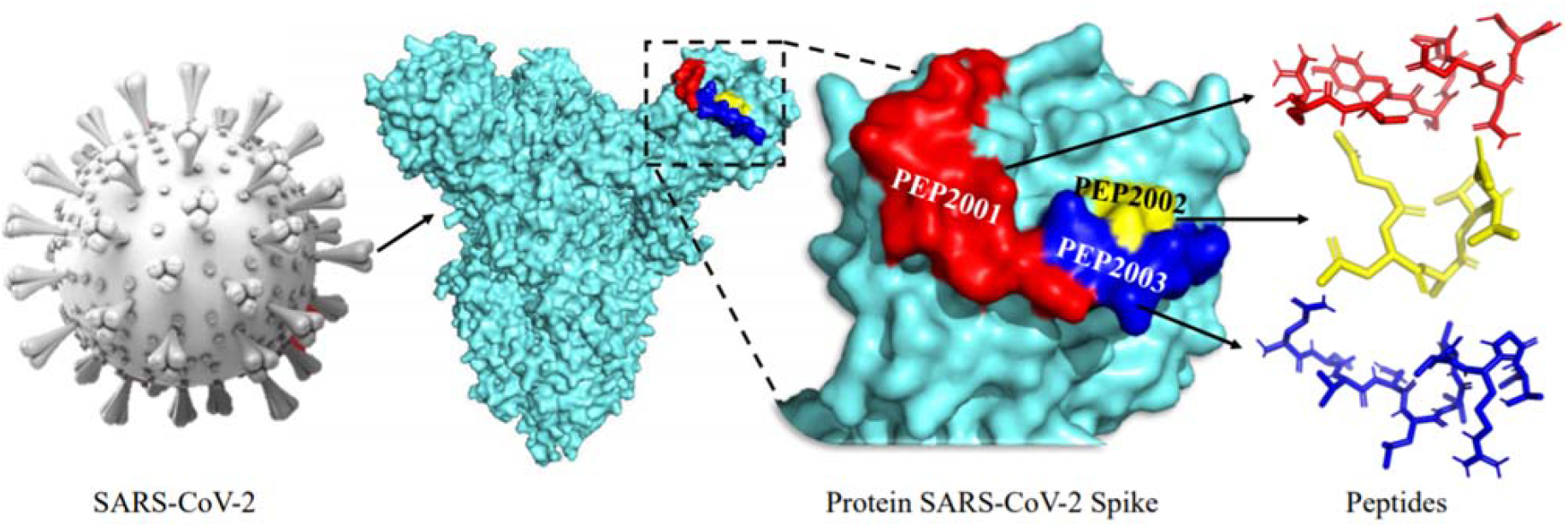
Structural models of peptides (A) PSPD2001, (B) PSPD2002, and (C) PSPD2003 that were synthesized in the present study.

**Table 1.** General information on the synthesis of the peptides used in the present study.

| Sequences | Codes | M.W. | Resins | Cleavage Solution |
| --- | --- | --- | --- | --- |
| Arg-Val-Tyr-Ser-Ser-Ala-Asn-Asn-Cys-COOH | PSPD2001 | 1013,09 | Fmoc-Cys- | 94% TFA |
|  |  |  | Wang | 2,5% H<sub>2</sub>O |
|  |  |  | SG*: 0,55 | 2,5% DODT* |
|  |  |  | mmol/g | 1% TIS* |
| Gln-Cys-Val-Asn-Leu-Thr-Thr-Arg-Thr-COOH | PSPD2002 | 1035,18 | Fmoc-Thr- | 94% TFA |
|  |  |  | Wang | 2,5% H<sub>2</sub>O |
|  |  |  | SG: 0,55 | 2,5% DODT |
|  |  |  | mmol/g | 1% TIS |
| Asn-Asn-Ala-Thr-Asn-COOH | PSPD2003 | 532,51 | Fmoc-Asn- | 95% TFA |
|  |  |  | Wang | 2,5% H<sub>2</sub>O |
|  |  |  | SG: 0,51 | 2,5% TIS |
|  |  |  | mmol/g |  |

M.W.: molecular weight; SG: substitution grade– Number of active sites available for the growth of the peptide chain. DODT: 2,2'-(Ethylenedioxy) diethanol; TIS: triisopropylsilane; TFA: trifluoroacetic acid.

**Table 2.**
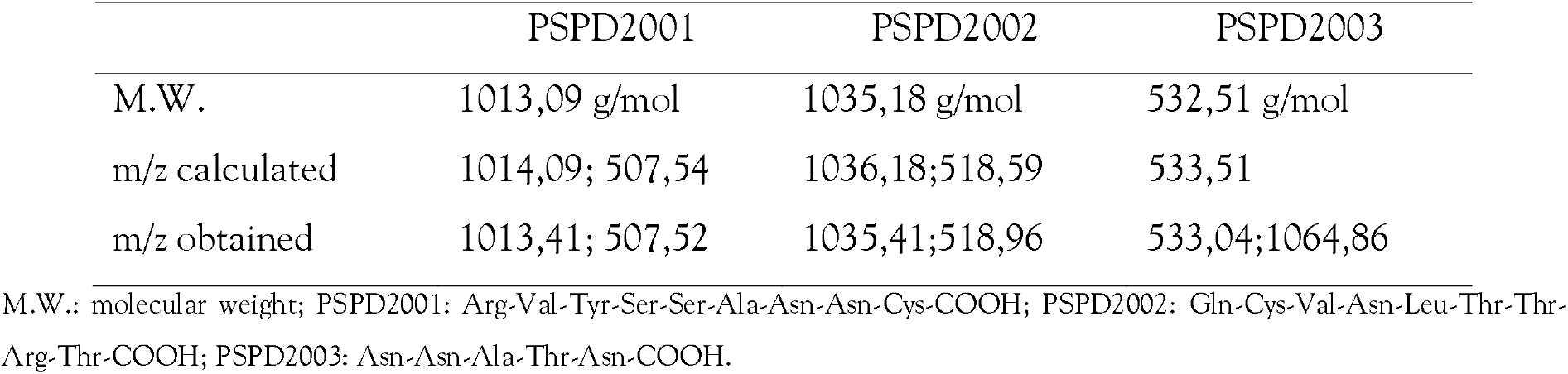
Mass spectrometry analysis after peptide purification

Regarding the alignment of the obtained sequences, our analyzes revealed the existence of similarities between the nucleotide sequences of the synthesized peptides and different regions conserved in SARS-Cov-2, whose comparisons were made from three datasets COVID, originated from Texas (USA), Iran, and Australia (Figure 2). These data demonstrated that the peptides PSPD2001, PSPD2002, and PSPD2003 are, in fact, part of the protein structure of the etiological agent of COVID-19. However, the sequences obtained for P. cuvieri (taxid: 218685) showed two main agreements belonging to five of the total of nine peptides found. The identification of possible linear epitopes was performed by BLAST with the Swissprot protein sequence database restricted to P. cuvieri. All peptides obtained in the sequencing were evaluated. However, only the main results found for five peptides are being presented in Table S1, two in the form SARS-CoV-2, in addition to the analysis made for the consensus obtained in alignment with Clustal W.

**Figure 2.**
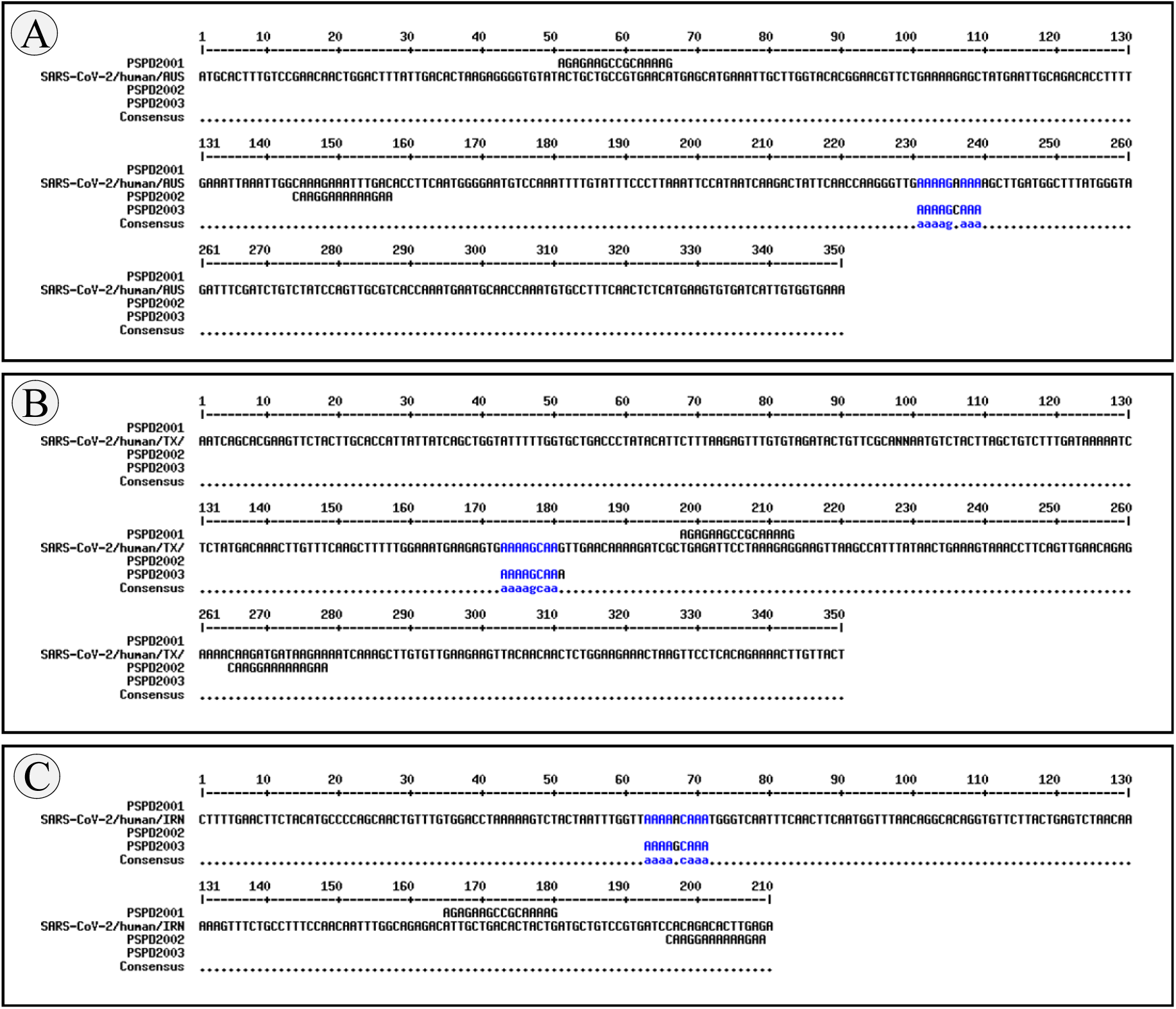
Alignment of the nucleotide sequence encoding PSPD2001, PSPD2002 and PSPD2003 with SARS-CoV-2 obtained from the (A) COVID dataset from Australia (SARS-Cov-2 / human / AUS), (B) Texas, USA (SARS-Cov-2 / human / TX) and (C) Iran (SARS-Cov-2 / human / IRN). The blue markings refer to similar nucleotides between the synthesized peptides and those present in SARS-CoV-2.

The in vivo experiments revealed that short exposure to SARS-CoV-2 Spike peptides was able to induce significant biochemical changes in P. cuvieri tadpoles. After 24 h of exposure, we observed that the peptides PSPD2002 and PSPD2003 (100 and 500 ng/mL) induced a significant increase in nitrite production (the indirect measurement of NO (Soneja et al. 2005) and hydrogen peroxide (Figure 4A-B, respectively), which in association with the higher levels of ROS (Figure 4D), suggest an increase in oxidative stress processes in the animals. The PSPD2003 peptide, in particular, demonstrated an even more significant effect on NO production, exceeding a 30% increase, to the control group, in both tested concentrations (100 and 500 ng/ml); almost 60% increase in hydrogen peroxide levels in the group exposed to 500 ng/mL, and 29% ROS in the animals treated with 100 ng/mL. However, we did not observe significant differences between the groups regarding MDA levels (Figure S2-A), suggesting that the treatments did not intensify the lipid peroxidation processes in the tadpoles.

**Figure 3.**
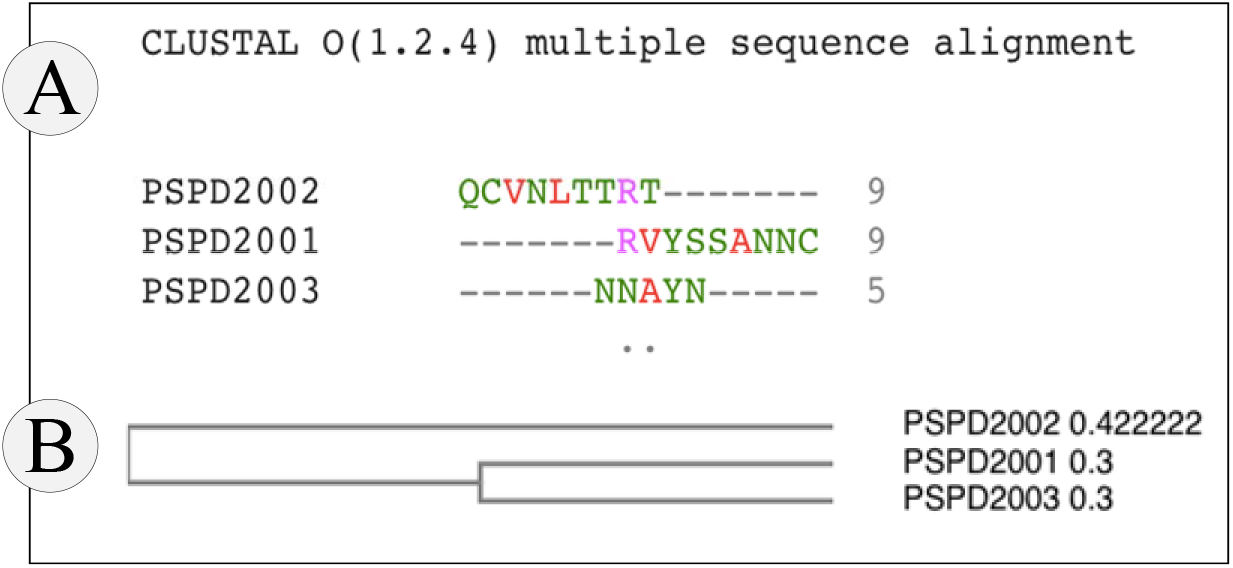
(A) Peptide alignment and (B) Guide Tree Phylogram. Both for the Physalaemus cuvieri form (taxid: 218685) by the Clustal W program. The green regions highlight 50% of the conserved region and, in red, 50% to 85% of the conserved region.

**Figure 4.**
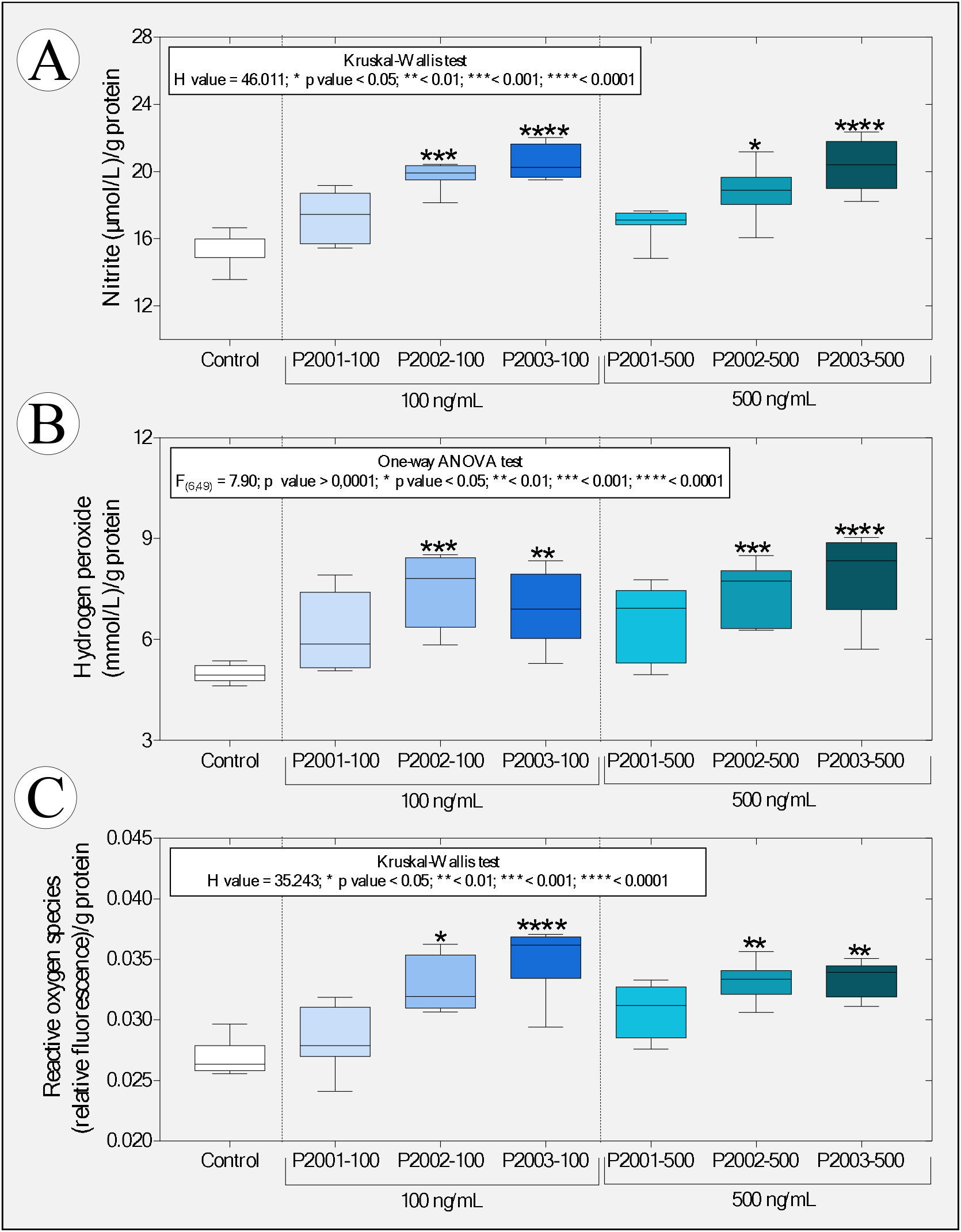
Boxplot of data obtained from predictive oxidative stress biomarkers [(A) nitrite levels, (B) hydrogen peroxide and (C) reactive oxygen species] in tadpoles of P. cuvieri (phase 27G) exposed or not to peptides PSPD 2001, 2002 and 2003 of the SARS-CoV-2 Spike protein. The summaries of the statistical analyzes are shown in the upper left corner of the graphs. Asterisks indicate significant differences between the respective groups and the control group. (n = 50 animals/group).PSPD2001: Arg-Val-Tyr-Ser-Ser-Ala-Asn-Asn-Cys-COOH; PSPD2002: Gln-Cys-Val-Asn-Leu-Thr-Thr-Arg-Thr-COOH; PSPD2003: Asn-Asn-Ala-Thr-Asn-COOH.

Similar to the previous findings, we observed that the animals exposed to PSPD2002 and PSPD2003 showed an increase, in a concentration-dependent manner, of the activity of the enzymes SOD and catalase (Figure 5A-B), with these data being positively and significantly correlated with the increase in the levels of nitrite, peroxide hydrogen and ROS (Figure 5C-D). We also observed that PSPD2003, once again, induced more intense effects on the antioxidant activity; there was an increase above 36% as compared to the control group for the two concentrations tested (100 and 500 ng/mL). The levels of SOD and catalase in the tadpoles exposed to PSPD2002 fragments were 28.9% higher than those reported in the control group. However, the SOD/catalase ratio was unaffected or decreased which indicates the relative balance between hydrogen peroxide synthesis by SOD and its decomposition by catalase (Figure 2S-B).

**Figure 5.**
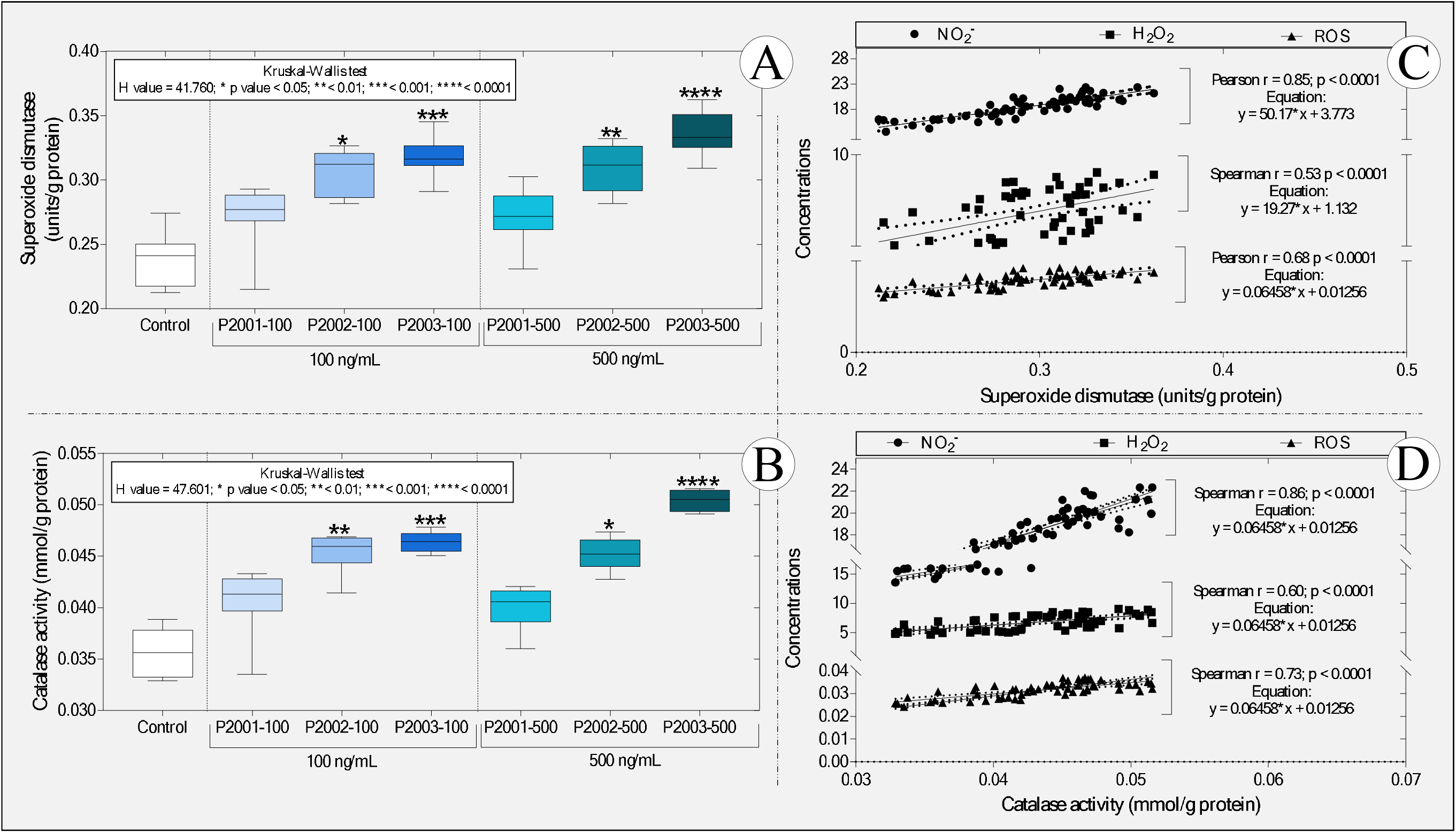
Boxplot of the activity of the enzymes (A) superoxide dismutase and (B) catalase, as well as correlations between the levels of (C) superoxide dismutase and (D) catalase and the different predictive biomarkers of oxidative stress. NO2-: nitrite; H2O2: hydrogen peroxide and ROS: reactive oxygen species. In “A” and “B,” the statistical analyses’ summaries are shown in the graphs’ upper left corner. Asterisks indicate significant differences between the respective groups and the control group. (n = 50 animals / group). PSPD2001: Arg-Val-Tyr-Ser-Ser-Ala-Asn-Asn-Cys-COOH; PSPD2002: Gln-Cys-Val-Asn-Leu-Thr-Thr-Arg-Thr-COOH; PSPD2003: Asn-Asn-Ala-Thr-Asn-COOH.

These data are exciting since they corroborate previous studies that describe the critical role the SARS-CoV-2 Spike protein in inducing oxidative stress in COVID-19 infection [see the review of Suhail et al. (2020)] while demonstrating that the peptides evaluated, even in a non-host organism, can cause metabolic disorders related to the increase in reactive species. On the other hand, the impairment of antioxidant defenses observed in several viral infections (Fraternale et al., 2006), including COVID-19 (Baradaran et al., 2020; Polonikov, 2020; Bayindir & Bayindir, 2020; Abouhashem et al., 2020), was not evident in the studied organism. These data also reinforce the hypothesis that the responses to the peptide fragments tested may be different between hosts and non-hosts of SARS-CoV-2; they also confirm the ability of peptides PSPD2002 and PSPD2003 to induce metabolic changes that alter REDOX homeostasis towards oxidative stress in tadpoles.

The proportional increase in oxidative stress biomarkers and the activity of SOD and catalase enzymes [two essential and indispensable molecules in cellular antioxidant defense strategies -Nishikawa et al. (2009), Hu & Tirelli (2012) and Ighodaro & Akinloye (2018)] reinforces our hypothesis, showing that the increase in antioxidant defenses does not seem to have been sufficient to reduce oxidative stress. The proposition of an action mechanism explaining the increase in these enzymes’ activity is very incipient, either due to our study’s pioneering nature or the need to deepen biochemical assessments in future studies. However, it is tempting to speculate that the interactions between PSPD2002 and PSPD2003 peptides and antioxidant enzymes evaluated in tadpoles (confirmed by molecular docking) have induced functional changes in SOD and catalase, similarly to what was observed by Jing et al. (2020) by exposing hepatocytes isolated from C57BL6 mice to different concentrations of naphthalene. The data obtained from the molecular docking reinforces our hypothesis by confirming the affinity between the PSPD2002 and PSPD2003 peptides and the referred enzymes and the existence of interactions with residues from all tested moorings (Figure 6). In the interactions with PSPD2002, it was possible to verify several hydrogen bonds in the threonine mixture (T9), revealing the potencies of the binding affinities and the central region of interaction in the active sites of the tested targets. In contrast, PSPD2003 interactions showed ?20 hydrogen interactions in all the tested couplings, with the structures of valine (V2) and serine (S4) (central of the peptide) considered to have the best affinity region of the ligand.

**Figure 6.**
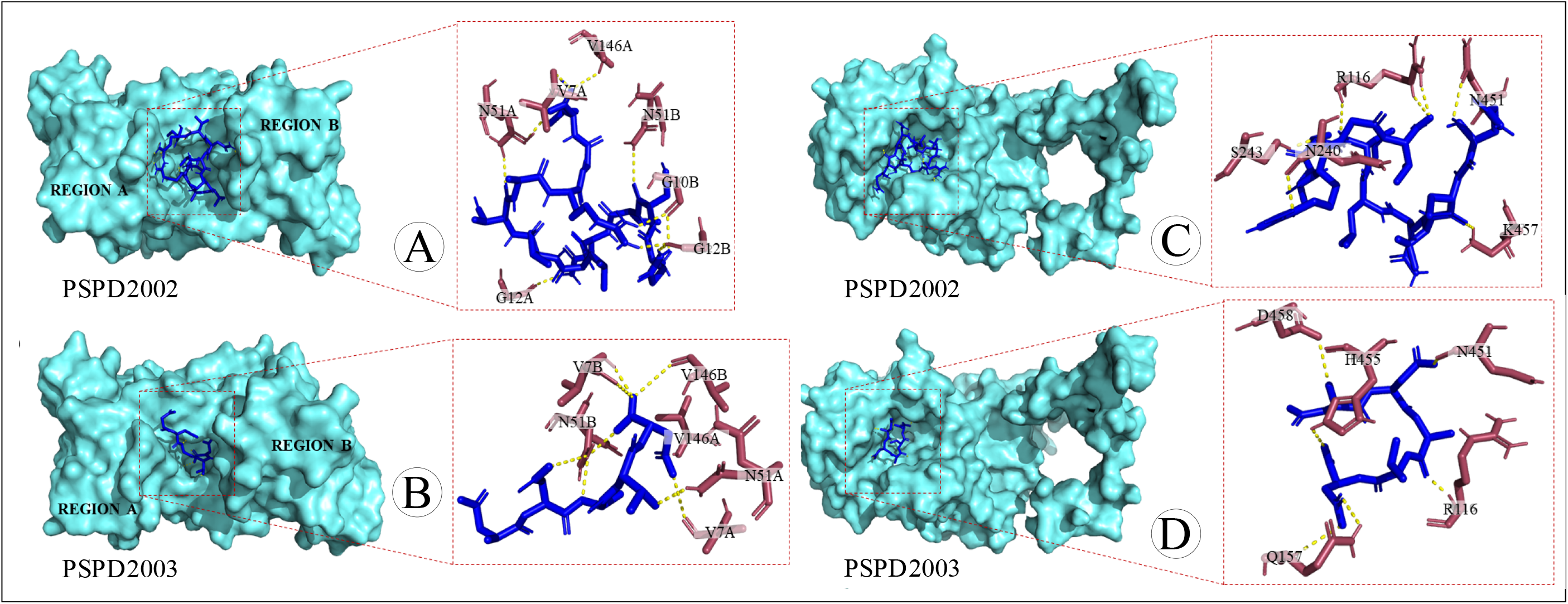
Three-dimensional surface-ligand coupling of interactions between peptides PSPD2002 and PSPD-2003 and the enzyme (A-B) superoxide dismutase (SOD) and (C-D) catalase (C-D), all in surface mode and highlighted active site. In “B and D”, we also observe regions A and B of the homo-dimer structure. Interaction residues in “A” (SOD-PSPD2002): G12A; N51A *; V7A; V146B; G10B *; G12B *; N51B (affinity (kcal / mol) = −8.3). In “B” (SOD-PSPD2002): N51A; V7A; V146A; N51B **; V7B; V146B * (affinity (kcal / mol) = −8.6). In “C” (Catalase-PSPD2002): K457; N240; N451; R116 **; S243 (affinity (kcal / mol) = −9.3). In “D” (Catalase-PSPD2003): D458; H455; N451; Q157 *; R116 (affinity (kcal / mol) = – 6.8). An “asterisk” indicates two interactions in the same residue. Two “asterics” indicate the existence of three interactions in the same residue.

An increase in NO production (inferred by high levels of nitrite) in tadpoles exposed to PSPD2002 and PSPD2003 (Figure 4A) suggests that the production of this free radial gas constitutes a standard response to the constituents of the SARS-CoV protein −2 Spike of the new coronavirus, both in the evaluated non-host organism and in humans infected with SARS-CoV-2. The ability of NO (both endogenous and exogenous) to inhibit the replication cycle of other viruses in the Coronaviridae family, affecting their proteins and reducing viral RNA (Chen et al., 2004; Keyaerts et al., 2004; Åkerström et al., 2005; Jung et al., 2010), has even motivated studies, whose preliminary results point to its potential therapeutic use in patients infected with SARS-CoV-2 (Alvarez et al., 2020). Alternatively, we cannot neglect the hypothesis of increased NO in tadpoles due tothe innate immune response modulated by the peptides, with a consequent increase in the production of inflammatory cytokines. In this case, studies reporting a positive correlation between NO production and increased pro-inflammatory cytokine levels (TNF-α, IL-6, IL-17, IL-12, and interferon-γ) in patients with COVID-19 reinforce our hypothesis (Karki et al., 2020; Del Valle et al., 2020; Costela-Ruiz et al., 2020).

We also evaluated the peptides’ possible neurotoxicity in tadpoles exposed to the peptide fragments of the SARS-CoV-2 Spike protein. Interestingly, we observed that 100 ng/mL PSPD2003 induced an increase greater than 220% concerning the control group. However, at a concentration of 500 ng/mL, all the peptides evaluated exerted an effect in the cholinergic system, causing an increase in the activity of AChE (Figure 7). While the peptides PSPD2001 and PSPD2002 induced increases of 219 and 553.8% in relation to AChE activity in the control group’s animals, respectively; the PSPD2003 peptide impressively induced an even more significant increase (697.3%). Therefore, these data confirmed the initial hypothesis that the SARS-CoV-2 Spike fragments induce neurotoxic effects, inferred by the stimulatory effect of the cholinergic system of the animals evaluated, especially in those exposed to the highest concentration (500 ng/mL) of the peptides.

**Figure 7.**
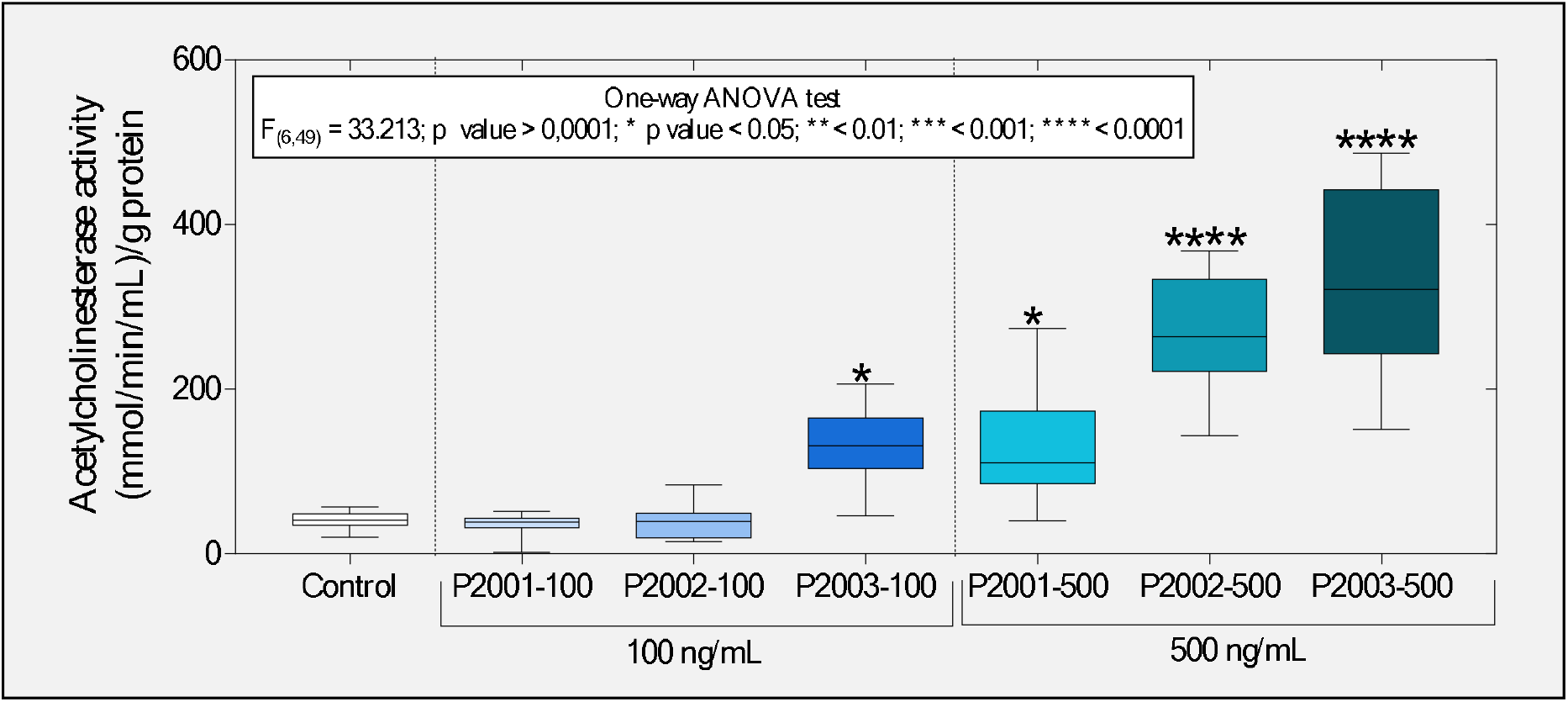
Boxplot of the enzyme acetylcholinesterase activity evaluated in tadpoles of P. cuvieri exposed or not to the peptides PSPD 2001, 2002, and 2003 of the SARS-CoV-2 Spike protein. The summaries of the statistical analyzes are shown in the upper left corner of the graphs. Asterisks indicate significant differences between the respective groups and the control group. (n = 50 animals / group). PSPD2001: Arg-Val-Tyr-Ser-Ser-Ala-Asn-Asn-Cys-COOH; PSPD2002: Gln-Cys-Val-Asn-Leu-Thr-Thr-Arg-Thr-COOH; PSPD2003: Asn-Asn-Ala-Thr-Asn-COOH.

Interestingly, these data differ from other studies that report suppression in AChE induced by increased cellular oxidative stress (Flora et al., 2013; Kayode et al., 2016; Bali et al., 2019; Ezeoyili et al., 2019; Pala et al., 2019; Ibrahim et al., 2020). In general, such studies argue that this can occur due to the deterioration of neurotransmission and oxidative damage. Additionally, AChE inhibition impairs oxidative phosphorylation and is followed by neuronal Ca^2+^influx and activation of nNOS, associated with the neurons’oxidative and nitrosative injury (Milatovic et al., 2006). However, the increased AChE activity observed in tadpoles exposed to the peptides may be related to the activation of the cholinergic anti-inflammatory pathway (CAP), which has been found beneficial in preventing inflammatory conditions such as sepsis and acute respiratory distress syndrome in animal models [see the review of Liu et al. (2020)]. As discussed by Osman (2020), CAP constitutes a neural mechanism that modulates inflammation through the release of acetylcholine (ACh), that have led to increased AChE synthesis to decompose higher levels of this neurotransmitter [see details in Tracey (2007)]. This mechanism has been reported in different studies involving patients infected with the new coronavirus (Bonaz et al., 2020; Mazloom et al., 2020; Pomara et al., 2020), strengthening the presumption that this mechanism may constitute another similar physiological response between SARS-CoV-2 non-host and host organisms. Besides, it is plausible to assume not only that the peptide composition of the SARS-CoV-2 Spike protein participates in the CAP activation (both in humans and in the evaluated tadpoles) but also that the neuroimmune system of the tadpoles has an essential role in responding to exposure of peptides PSPD2001, PSPD2002 and PSPD2003.

Alternatively, the tadpoles’ cholinergic system’s stimulation may also be explained by the direct interactions between the tested peptides and AChE, whose affinity was demonstrated in the molecular docking analysis (Figure 8). In this case, future studies will be useful to understand if these interactions induced a significant change in the association and catalysis mechanism or expansion of the enzyme efficiency with an increase of the substrate affinity to the active site (increasing the catalytic constant was increased and decreasing the Michaelis constant). In both situations, a significant increase in AChE activity can occur, either as part of a compensatory mechanism that will aim to compensate for the enzyme’s catalytic deficit or as a more efficient response to the increased release of acetylcholine in synaptic clefts via CAP activation. The hypothesis that increased AChE activity in these animals was associated with positive AChE gene regulation due to Spike protein peptides’ inhibitory effect needs to be tested in future studies.

**Figure 8.**
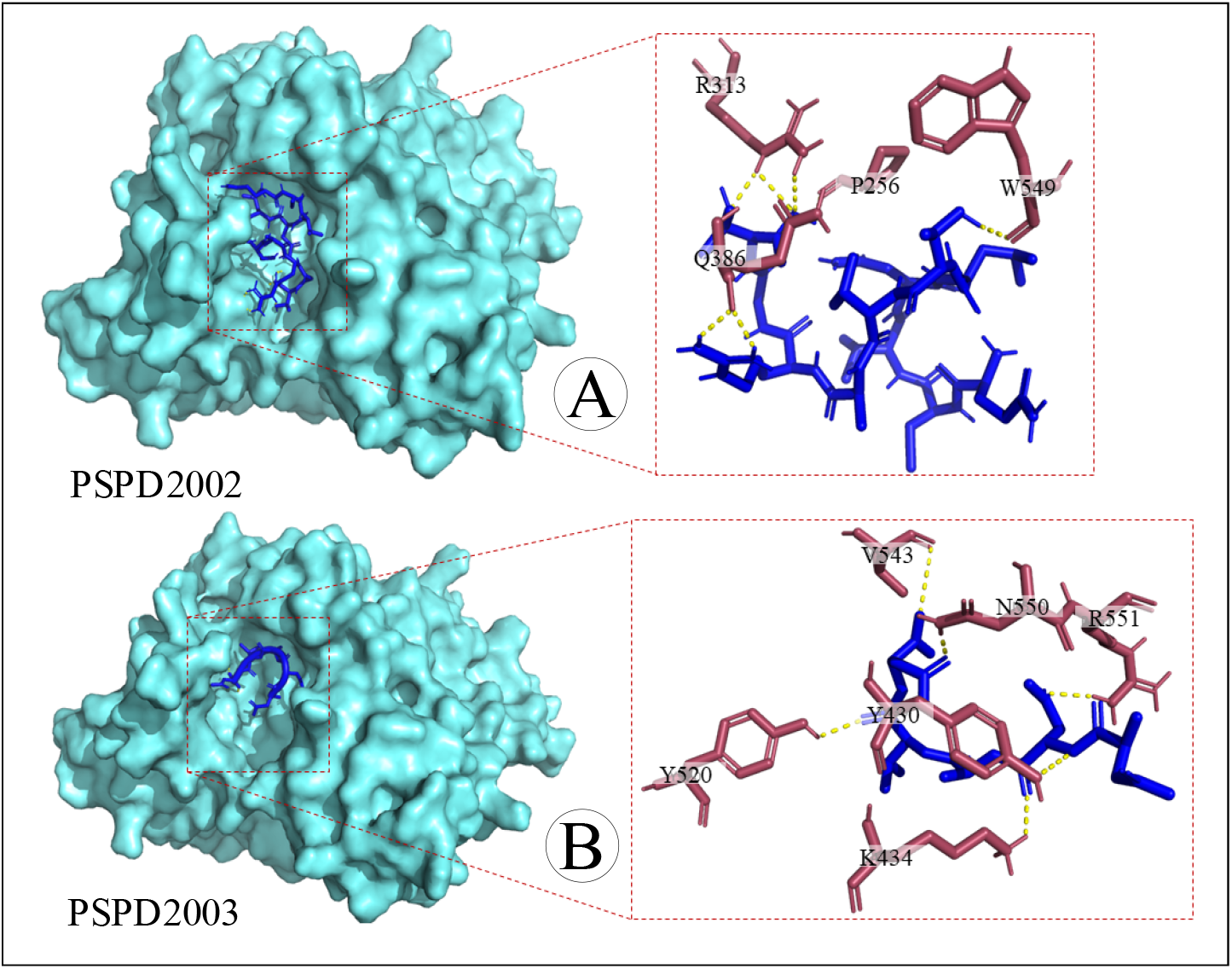
Three-dimensional surface-ligand coupling of interactions between peptides (A) PSPD2002 and (B) PSPD-2003 and the enzyme acetylcholinesterase, all in surface mode and highlighted active site. Interaction residues in “A” (AChE-PSPD2002): P256; Q386 *; R313 **; W549 (affinity (kcal / mol) = −9.4). In “B” (AChE-PSPD2003): K434; N550; R551; V543; Y430; Y520 (affinity (kcal / mol) = −8.4). An “asterisk” indicates two interactions in the same residue. Two “asterics” indicate the existence of three interactions in the same residue.

Finally, it is essential to emphasize that although our study gathers clear and pioneering evidence on the negative impact of the SARS-CoV-2 Spike fragments (especially PSPD2002 and PSPD2003) on the biochemical parameters evaluated in P. cuvieri tadpoles, many questions about the consequences of the presence of these fragments in the aquatic environment remains obscure. The evaluation of the effects of prolonged exposure to the tested peptides (in higher and lower concentrations), the use of other experimental models (expanding the environmental representativeness), and the use of multiple toxicity biomarkers are some future investigative perspectives. Equally important will be to deepen the mechanisms of action of the peptides of the SARS-CoV-2 Spike protein when in direct contact with non-host organisms of the new coronavirus. Approaches of this nature can significantly expand our knowledge of the impact of COVID-19 on the environment and the functioning of ecosystems, and support the proposal for strategies to remedy or mitigate aquatic contamination by SARS-Cov2 particles.

## 4. CONCLUSIONS

From a systemic approach that included the synthesis, cleavage, purification, and alignment of peptides to P. cuvieri tadpoles’ exposure to peptide fragments of Spike protein, we gathered evidence that confirms the toxicity of viral constituents in the evaluated animal model. The increase in predictive biomarkers of REDOX imbalance and neurotoxic action is, therefore, an insight into how aquatic particle contamination of SARS-CoV-2 can constitute additional environmental damage to the COVID-19 pandemic. In this sense, we strongly suggest conducting further studies necessary to understand the real magnitude of the biological/environmental impact of COVID-19.

## 5. ACKNOWLEDGMENT

This work was supported by São Paulo Research Foundation (FAPESP 2020/05761-3), Brazilian National Research Council (CNPq) (426531/2018-3) and Instituto Federal Goiano for the financial support (23219.001309.2020-19). Malafaia G. holds productivity scholarship from CNPq (307743/2018-7).

## SUPPLEMENTARY MATERIAL

**Table S1.** Summary information on the methodological procedures adopted in the purification stage of the present study’s compounds.

| Compound | Retention time (min) | Purificationmethod |
| --- | --- | --- |
| PSPD2001 | 7,7 | Gradient 3 to 43% B in 90 min |
| PSPD2002 | 10,0 | Gradient 10 to 50% B in 90 min |
| PSPD2003 | 2,0 e 13,3 | Isocratic 2% to B in 20 min and gradient 2 to 53% B in 90 min |
| PSPD2001: Arg-Val-Tyr-Ser-Ser-Ala-Asn-Asn-Cys- COOH; PSPD2002: Gln-Cys-Val-Asn-Leu-Thr-Thr-Arg-Thr-COOH; PSPD2003: Asn-Asn-Ala-Thr-Asn-COOH. |  |  |

**Figure S1.**
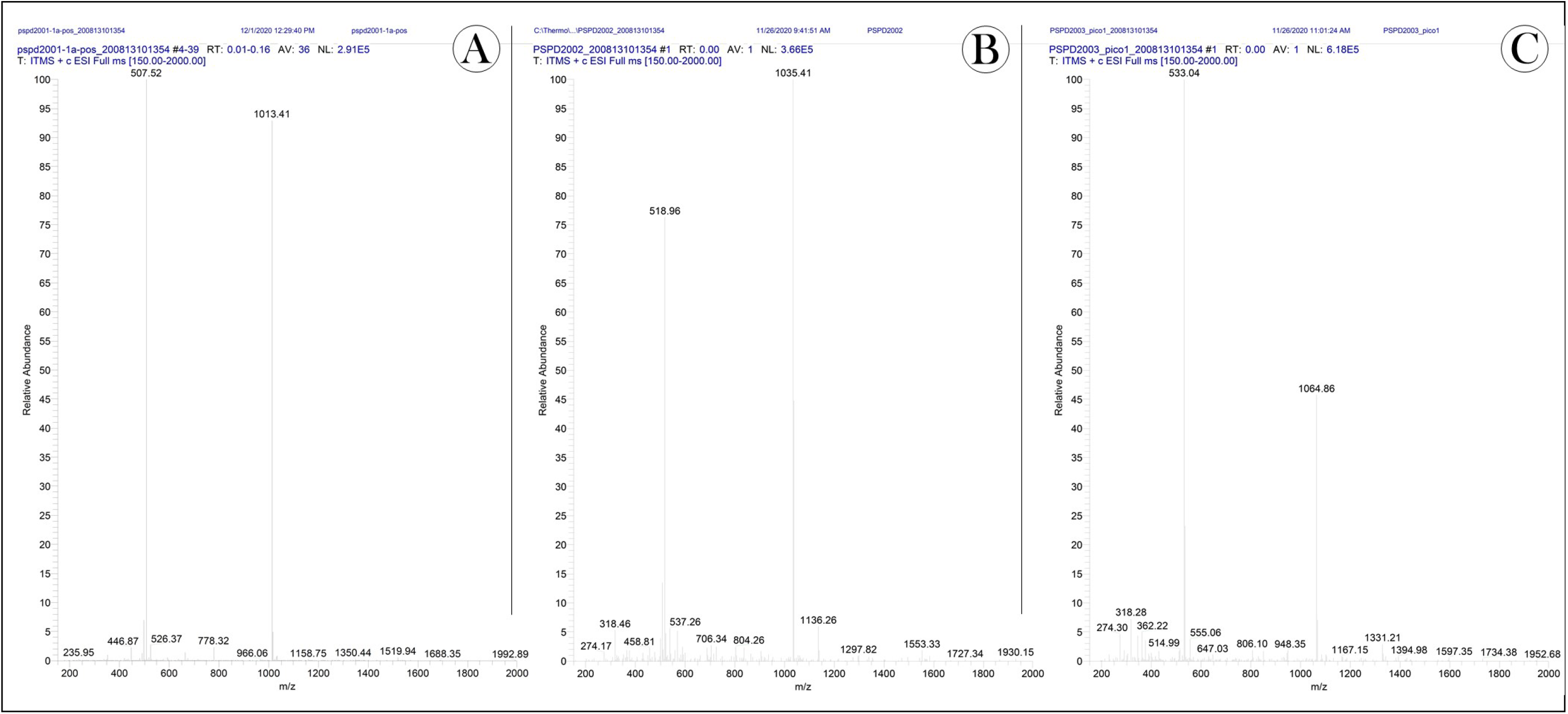
Mass spectra obtained for peptides (A) PSPD2001, (B) PSPD2002 and (C) PSPD2003. PSPD2001: Arg-Val-Tyr-Ser-Ser-Ala-Asn-Asn-Cys-COOH; PSPD2002: Gln-Cys-Val-Asn-Leu-Thr-Thr-Arg-Thr-COOH; PSPD2003: Asn-Asn-Ala-Thr-Asn-COOH.

**Figure S2.**
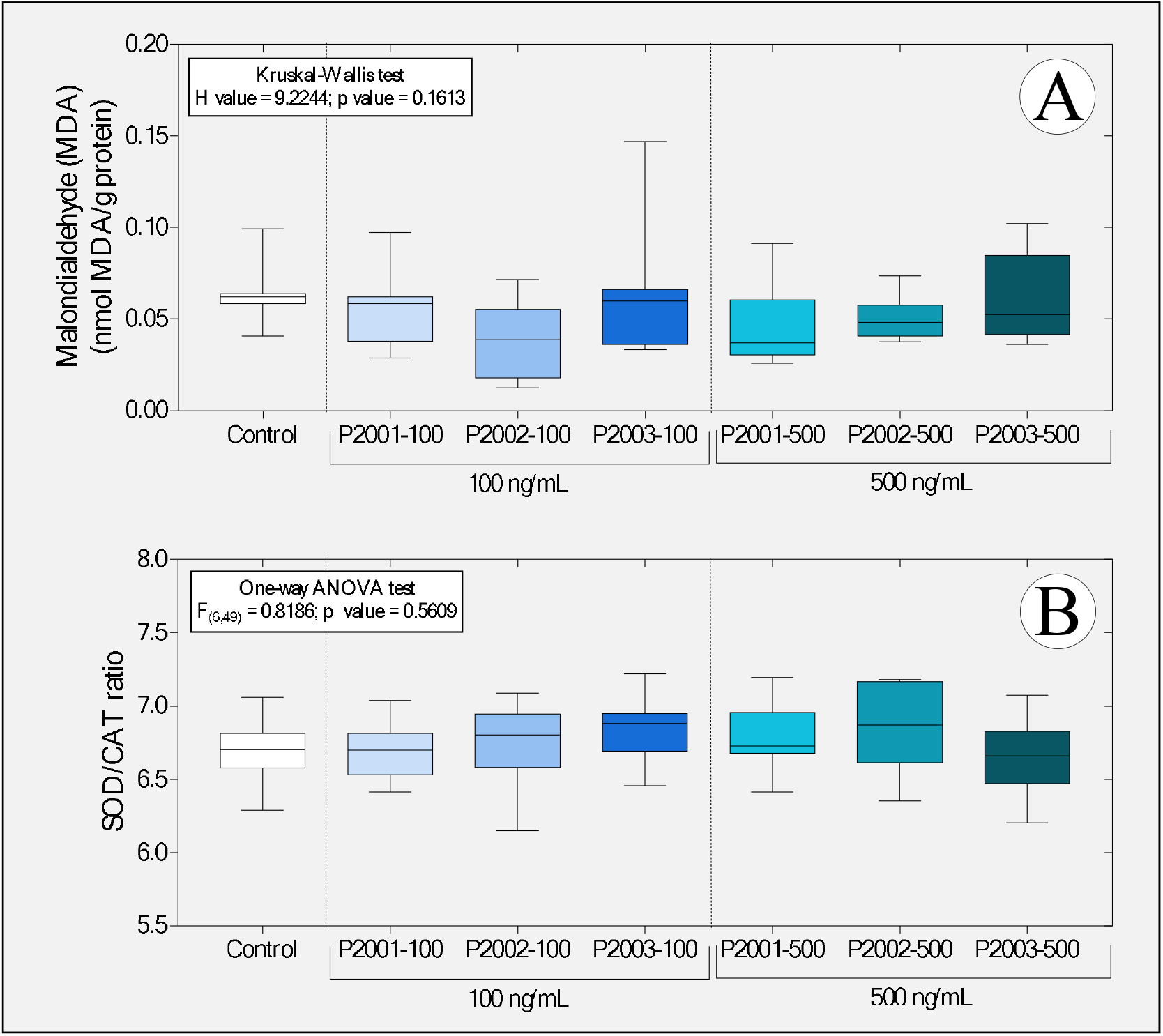
Boxplot of (A) concentrations of malondialdehyde and (B) SOD / CAT ratio in tadpoles of P. cuvieri (phase 27G) exposed or not to PSPD peptides 2001, 2002, and 2003 of the SARS-CoV-2 Spike protein. Summaries of statistical analyzes are shown in the upper left corner of the graph. (n = 50 animals / group). PSPD2001: Arg-Val-Tyr-Ser-Ser-Ala-Asn-Asn-Cys-COOH; PSPD2002: Gln-Cys-Val-Asn-Leu-Thr-Thr-Arg-Thr-COOH; PSPD2003: Asn-Asn-Ala-Thr-Asn-COOH. SOD: superoxide dismutase; CAT: catalase.

**Table S1:**
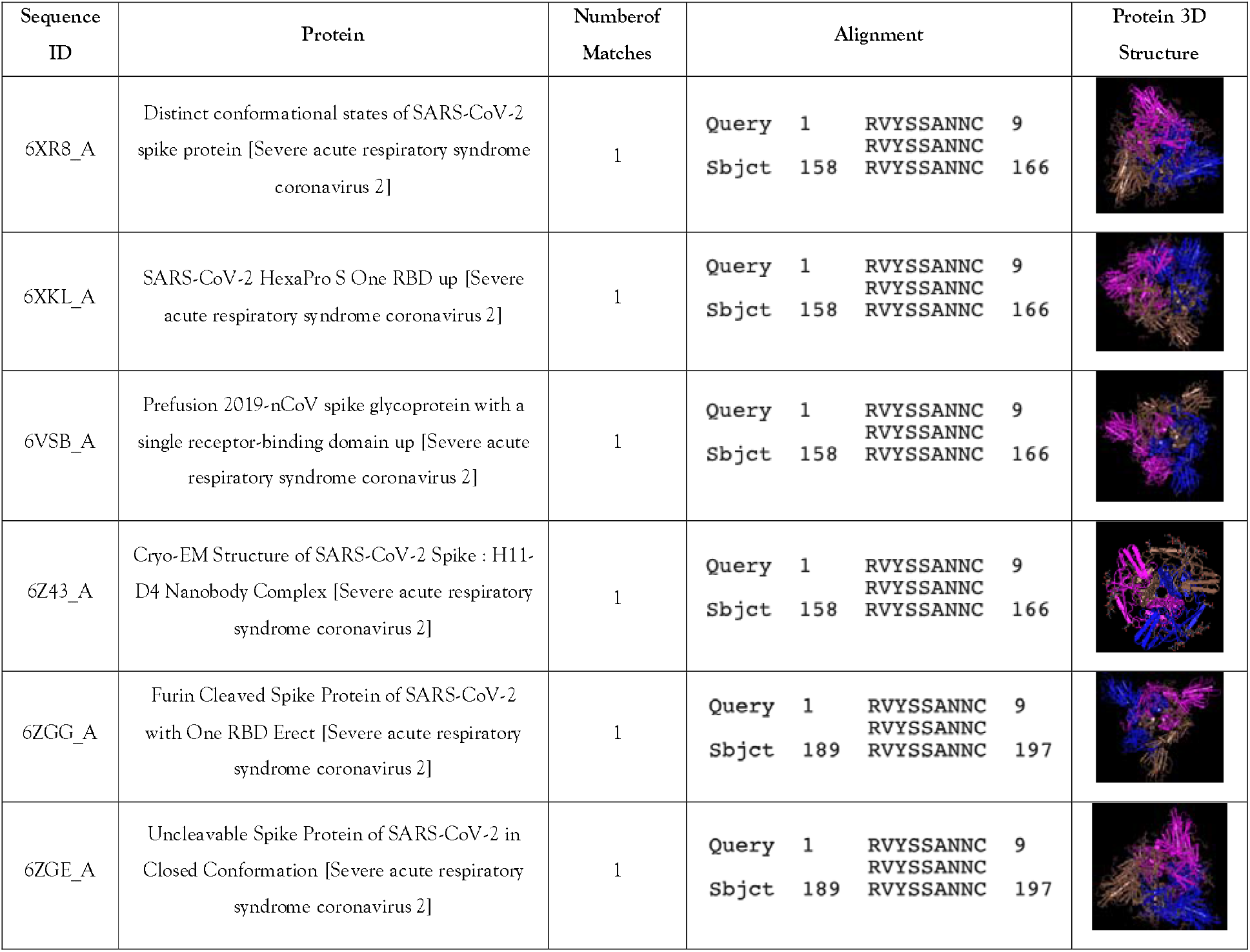
Alignment of the sequences and consensus for PSPD2001, showing the similarities found with the proteins noted in the GeneBank database by BLAST, with the Sequence ID, the protein’s name, the Number of Matches, the alignment and the Protein 3D Structure.

## Notes

### Competing Interest Statement

The authors have declared no competing interest.

## REFERENCES

Abouhashem, A. S., Singh, K., Azzazy, H. M., & Sen, C. K. (2020). Is Low Alveolar Type II Cell SOD3 in theLungsofElderlyLinkedtotheObservedSeverity of COVID-19?. Antioxidants & Redox Signaling.

Abu-Qdais, H. A., Al-Ghazo, M. A., & Al-Ghazo, E. M. (2020). Statistical analysis and characteristics of hospital medical waste under novel Coronavirus outbreak. Global Journal of Environmental Science and Management, 6(Special Issue (Covid-19)), 21–30.

Abu-Rayash, A., & Dincer, I. (2020). Analysis of the electricity demand trends amidst the COVID-19 coronavirus pandemic. Energy Research & Social Science, 68, 101682.

Adrieli Sachett, Matheus Gallas-Lopes, Greicy M MConterato, Radharani Benvenutti, Ana P Herrmann, Angelo Piato 2020. Quantification of thiobarbituric acid reactive species (TBARS) optimized for zebrafish brain tissue. protocols.io. https://dx.doi.org/10.17504/protocols.io.bjp8kmrw

Åkerström, S., Gunalan, V., Keng, C. T., Tan, Y. J., & Mirazimi, A. (2009). Dual effectofnitric oxide on SARS-CoVreplication: viral RNA production and palmitoylationofthe S protein are affected. Virology, 395(1), 1–9.

Åkerström, S., Mousavi-Jazi, M., Klingström, J., Leijon, M., Lundkvist, Å., & Mirazimi, A. (2005). Nitric oxide inhibitsthereplicationcycleofsevereacuterespiratorysyndrome coronavirus. Journal of virology, 79(3), 1966–1969.

Alvarez RA, Berra L, Gladwin MT. Home nitric oxide therapyfor COVID-19. Am J RespirCritCareMed, 202(1): 16–20, 2020.

Araújo, A. P. C., de Melo, N. F. S., de Oliveira Junior, A. G., Rodrigues, F. P., Fernandes, T., de Andrade Vieira, J. E., … & Malafaia, G. (2020a). How much are microplastics harmful to the health of amphibians? A study with pristine polyethylene microplastics and Physalaemus cuvieri. Journal of hazardous materials, 382, 121066.

Araújo, A. P. C., Gomes, A. R., & Malafaia, G. (2020b). Hepatotoxicity of pristine polyethylene microplastics in neotropical physalaemus cuvieri tadpoles (Fitzinger, 1826). Journal of Hazardous Materials, 386, 121992.

Bali, Y. A., Kaikai, N. E., Ba-M’hamed, S., & Bennis, M. (2019). Learning and memoryimpairmentsassociatedtoacetylcholinesteraseinhibition and oxidative stress followingglyphosatebased-herbicideexposure in mice. Toxicology, 415, 18–25.

Bangaru, S., Ozorowski, G., Turner, H. L., Antanasijevic, A., Huang, D., Wang, X., … & Patel, N. (2020). Structural analysis of full-length SARS-CoV-2 spike protein from an advanced vaccine candidate. Science, 370(6520), 1089–1094.

Baradaran, A., Ebrahimzadeh, M. H., Baradaran, A., & Kachooei, A. R. (2020). Prevalenceofcomorbidities in COVID-19 patients: A systematicreview and meta-analysis. Archives ofBone and JointSurgery, 8(Suppl 1), 247.

Bayindir, M., & Bayindir, E. E. (2020). Synergic viral-bacterialco-infection in catalase-deficient COVID-19 patients causes suppressedinnateimmunity and lungdamagesduetodetrimentalelevationofhydrogenperoxideconcentration. Available at SSRN 3648292.

Behrendt, R., White, P., & Offer, J. (2016). Advances in Fmoc solid^phase peptide synthesis. Journal of Peptide Science, 22(1), 4–27.

Bivins, A., Greaves, J., Fischer, R., Yinda, K. C., Ahmed, W., Kitajima, M., … & Bibby, K. (2020). Persistence of SARS-CoV-2 in water and wastewater. Environmental Science & Technology Letters.

Blaustein, A. R., & Kiesecker, J. M. (2002). Complexity in conservation: lessons from the global decline of amphibian populations. Ecology letters, 5(4), 597–608.

Bonaz, B., Sinniger, V., & Pellissier, S. (2020). Targetingthecholinergicanti-inflammatorypathwaywithvagusnervestimulation in patientswith Covid-19?. Bioelectronic medicine, 6(1), 1–7.

Braz HLB, Silveira JAM, Marinho AD, de Moraes MEA, Moraes Filho MO, Monteiro HSA, Jorge RJB. In silico study of azithromycin, chloroquine and hydroxychloroquine and their potential mechanisms of action against SARS-CoV-2 infection. Int J AntimicrobAgents. 2020 Sep;56(3):106119. doi:10.1016/j.ijantimicag.2020.106119. Epub 2020 Jul 30. PMID: 32738306; PMCID:PMC7390782.

Carvalho, M., Bandeira de Mello Delgado, D., de Lima, K. M., de Camargo Cancela, M., dos Siqueira, C. A., & de Souza, D. L. B. (2020). Effects of the COVID⒡19 pandemic on the Brazilian electricity consumption patterns. International Journal of Energy Research, e5877.

Chen L, Liu P, Gao H, Sun B, Chao D, et al. (2004) Inhalationofnitric oxide in thetreatmentofsevereacuterespiratorysyndrome: a rescue trial in Beijing. Clin InfectDis 39: 1531–1535.

Chen, Y., Chen, L., Deng, Q., Zhang, G., Wu, K., Ni, L., … & Yang, J. (2020). The presence of SARS□CoV□2 RNA in the feces of COVID□19 patients. Journal of medical virology.

Colston, J. T., Chandrasekar, B., & Freeman, G. L. (2002). A novel peroxide-inducedcalciumtransientregulates interleukin-6 expression in cardiac-derivedfibroblasts. Journal of BiologicalChemistry, 277(26), 23477–23483.

Costela-Ruiz, V. J., Illescas-Montes, R., Puerta-Puerta, J. M., Ruiz, C., & Melguizo-Rodríguez, L. (2020). SARS-CoV-2 infection: the role ofcytokines in COVID-19 disease. Cytokine & Growth Factor Reviews.

Coughlan, L. (2020). Snatching the Crown from SARS-CoV-2. Cell Host & Microbe, 28(3), 360–363.

Chakraborty, I & Prasenjit, M. COVID-19 outbreak: Migration, effects on society, global environment and prevention. Science of the Total Environment, 2020, 138882.

Del Valle, D. M., Kim-Schulze, S., Hsin-hui, H., Beckmann, N. D., Nirenberg, S., Wang, B., … & Marron, T. (2020). Aninflammatorycytokinesignaturehelpspredict COVID-19 severity and death. medRxiv.

Del-Maestro, R. F., & McDonald, W. (1985). Oxidative enzymes in tissue homogenates. Handbook of methods for oxygen radical research, 291–296.

Elnemma, E. M. (2004). Spectrophotometric determination of hydrogen peroxide by a hydroquinone-aniline system catalyzed by molybdate. Bulletin of the Korean Chemical Society, 25(1), 127–129.

Elsamadony, M., Fujii, M., Miura, T., & Watanabe, T. (2020). Possible transmission of viruses from contaminated human feces and sewage: Implications for SARS-CoV-2. Science of the Total Environment, 755, 142575.

Emmanuel DaanobaSunkari, Harriet MatekoKorboe, Mahamuda Abu, TefideKizildeniz,

Estrela, F. N., Guimarães, A. T. B., Silva, F. G., da Luz, T. M., Silva, A. M., Pereira, P. S., & Malafaia, G. (2021). Effects of polystyrene nanoplastics on Ctenopharyngodonidella (grass carp) after individual and combined exposure with zinc oxide nanoparticles. Journal of Hazardous Materials, 403, 123879.

European Centre for Disease Prevention and Control (ECDC). COVID-19 situation update for the EU/EEA and the UK, as of week 50 2020. Available in: https://www.ecdc.europa.eu/en/cases-2019-ncov-eueea. Acesson: 21 december 2020.

Editorital, The Lancet Respiratory Medicine, COVID-19 transmission—up in the air, The Lancet Respiratory Medicine, Volume 8, Issue 12, 2020, Page 1159, ISSN 2213-2600, https://doi.org/10.1016/S2213-2600(20)30514-2.

Ezeoyili, I. C., Mgbenka, B. O., Atama, C. I., Ngwu, G. I., Madu, J. C., & Nwani, C. D. (2019). Changes in BrainAcetylcholinesterase and Oxidative Stress Biomarkers in AfricanCatfishExposedtoCarbendazim. JournalofAquatic Animal Health, 31(4), 371–379.

Farsalinos, K., Niaura, R., Le Houezec, J., Barbouni, A., Tsatsakis, A., Kouretas, D., … & Poulas, K. (2020). Nicotine and SARS-CoV-2: COVID-19 may be a disease of the nicotinic cholinergic system. Toxicology Reports.

Fernandes, BH, Feitosa, N. M., Barbosa, A. P., Bomfim, C. G., Garnique, A. M. B., Gomes, F. I. F., … & Charlie-Silva, I. (2020). Zebrafish studies on the vaccine candidate to COVID-19, the Spike protein: Production of antibody and adverse reaction. https://doi.org/10.1101/2020.10.20.346262

Flora, S. J., Mehta, A., Satsangi, K., Kannan, G. M., & Gupta, M. (2003). Aluminum-inducedoxidative stress in ratbrain: response tocombinedadministrationofcitricacid and HEDTA. ComparativeBiochemistry and PhysiologyPart C: Toxicology & Pharmacology, 134(3), 319–328.

Fraternale A, Paoletti MF, Casabianca A, Oiry J, Clayette P, Vogel JU, Cinatl J, Jr, Palamara AT, Sgarbanti R, Garaci E, Millo E, Benatti U, Magnani M. Antiviral and immunomodulatorypropertiesof new pro-glutathione (GSH) molecules. CurrMedChem. 2006;13(15):1749–1755. doi:10.2174/092986706777452542

Frost DR. Amphibian Species of the World: an Online Reference. Version 6.0. Available in: http://research.amnh.org/vz/herpetology/amphibia/. Access on: 11 march. 2017.

Gaudin, Raphael; Goetz, Jacky G. Tracking Mechanisms of Viral Dissemination In Vivo. Trends in Cell Biology, 2021.

Gornall, A.G.; Bardawill, C.J.; David, M. M. Determination of serum proteins by means of the biuret reaction. J. Biol. Chem. v.177, p.751–766, 1949.

Gosner KL. A simplified table for staging anuran embryos and larvae with notes on identification. Herpetologica, 16:183–190 1960.

Grant, E. H. C., Miller, D. A., & Muths, E. (2020). A Synthesis of Evidence of Drivers of Amphibian Declines. Herpetologica.

Graham, Katherine E., et al. SARS-CoV-2 RNA in Wastewater Settled Solids Is Associated with COVID-19 Cases in a Large Urban Sewershed. Environmental science & technology, 2020.

Guerrero-Latorre, L., Ballesteros, I., Villacrés-Granda, I., Granda, M. G., Freire-Paspuel, B., & Ríos-Touma, B. (2020). SARS-CoV-2 in river water: Implications in low sanitation countries. Science of the Total environment, 743, 140832.

Guimarães, A. T. B., Charlie-Silva, I., & Malafaia, G. (2020). TOXIC EFFECTS OF NATURALLY-AGED MICROPLASTICS ON ZEBRAFISH JUVENILES: A MORE REALISTIC APPROACH TO PLASTIC POLLUTION IN FRESHWATER ECOSYSTEMS. JournalofHazardousMaterials, 124833.

Guimarães, A. T. B., de Lima Rodrigues, A. S., Pereira, P. S., Silva, F. G., & Malafaia, G. (2021). Toxicity of polystyrene nanoplastics in dragonfly larvae: An insight on how these pollutants can affect bentonic macroinvertebrates. Science of The Total Environment, 752, 141936.

Guy, C. A., & Fields, G. B. (1997). [5] Trifluoroacetic acid cleavage and deprotection of resinbound peptides following synthesis by Fmoc chemistry. Methods in enzymology, 289, 67–83.

Harrison, A. G., Lin, T., & Wang, P. (2020). Mechanisms of SARS-CoV-2 transmission and pathogenesis. Trends in immunology.

Henry, R. J.; Sobel, C.; Berkman, S. Interferences with biuret methods for serum proteins. Use of Benedict’s qualitative glucose reagent as a biuret reagent. Anal. Chem. v.29, p.1491–1495, 1957.

Herek, J. S., Vargas, L., Trindade, S. A. R., Rutkoski, C. F., Macagnan, N., Hartmann, P. A., & Hartmann, M. T. (2020). Can environmental concentrations of glyphosate affect survival and cause malformation in amphibians? Effects from a glyphosate-based herbicide on Physalaemus cuvieri and P. gracilis (Anura: Leptodactylidae). Environmental Science and Pollution Research, 1–12.

Higgins, D. G., Thompson, J. D., & Gibson, T. J. (1996). [22] Using CLUSTAL for multiple sequence alignments. In Methods in enzymology (Vol. 266, pp. 383–402). Academic Press.

Hu, P., & Tirelli, N. (2012). Scavenging ROS: superoxidedismutase/catalasemimeticsbythe use ofanoxidation-sensitivenanocarrier/enzymeconjugate. Bioconjugatechemistry, 23(3), 438–449.

Huang, Y., Yang, C., Xu, X. F., Xu, W., & Liu, S. W. (2020). Structural and functional properties of SARS-CoV-2 spike protein: potential antivirus drug development for COVID-19. Acta PharmacologicaSinica, 41(9), 1141–1149.

Ibrahim, K. A. E. M., Abdelrahman, S. M., Elhakim, H. K., & Ragab, E. A. (2020). Singleorcombinedexposuretochlorpyrifos and cypermethrinprovokeoxidative stress and downregulation in monoamine oxidase and acetylcholinesterase gene expressionoftherat’sbrain. EnvironmentalScience and PollutionResearch, 1–12.

Pomara, N., & Imbimbo, B. P. (2020). Impairmentofthecholinergicanti-inflammatorypathway in oldersubjectswithsevere COVID-19. Medical hypotheses.

Ighodaro, O. M., & Akinloye, O. A. (2018). First line defenceantioxidants-superoxidedismutase (SOD), catalase (CAT) and glutathioneperoxidase (GPX): Their fundamental role in theentireantioxidantdefencegrid. Alexandria journal of medicine, 54(4), 287–293.

Ighodaro, O. M., & Akinloye, O. A. (2018). First line defence antioxidants-superoxide dismutase (SOD), catalase (CAT) and glutathione peroxidase (GPX): Their fundamental role in the entire antioxidant defence grid. Alexandria journal of medicine, 54(4), 287–293.

Jetz, W, and R. A. Pyron. 2018. The interplay of past diversification and evolutionary isolation with present imperilment across the amphibian tree of life. Nature Ecology & Evolution 2:850–858.

Jing, M., Han, G., Wan, J., Zhang, S., Yang, J., Zong, W., … & Liu, R. (2020). Catalase and superoxidedismutase response and theunderlying molecular mechanismfornaphthalene. ScienceofThe Total Environment, 139567.

Jones, D. L., Baluja, M. Q., Graham, D. W., Corbishley, A., McDonald, J. E., Malham, S. K., … & Wilcox, M. H. (2020). Shedding of SARS-CoV-2 in feces and urine and its potential role in person-to-person transmission and the environment-based spread of COVID-19. Science of the Total Environment, 749, 141364.

Galindo-Villegas, Jorge. The zebrafish disease and drug screening model: A strong ally against Covid-19. Frontiers in Pharmacology, 2020, 11: 680.

Jung K, Gurnani A, Renukaradhya GJ, Saif LJ (2010) Nitric oxide iselicited and inhibits viral replication in pigsinfectedwithporcinerespiratory coronavirus butnotporcinereproductive and respiratorysyndrome virus. VetImmunolImmunopathol 136 335–339. [Crossref]

Kampf, G., Todt, D., Pfaender, S., & Steinmann, E. (2020). Persistence of coronaviruses on inanimate surfaces and their inactivation with biocidal agents. Journal of Hospital Infection, 104(3), 246–251.

Karki, R., Sharma, B. R., Tuladhar, S., Williams, E. P., Zalduondo, L., Samir, P., … & Schreiner, P. (2020). Synergismof TNF-α and IFN-γtriggersinflammatorycelldeath, tissuedamage, and mortality in SARS-CoV-2 infection and cytokine shock syndromes. Cell.

Kayode AO, Sulaiman O, Emmanuel AG, Dorcas W. Acetylcholinesteraseactivity and oxidative stress indices in cerebellum, cortex and hippocampusofratsexposedto lead and manganese; InternationaJournlofBiologicalResearch, 4(2): 157–164, 2016.

Keech, C., Albert, G., Cho, I., Robertson, A., Reed, P., Neal, S., … & Smith, G. (2020). Phase 1–2 trial of a SARS-CoV-2 recombinant spike protein nanoparticle vaccine. New England Journal of Medicine.

Keyaerts E, Vijgen L, Chen L, Maes P, Hedenstierna G (2004) Inhibitionof SARS-coronavirus infection in vitro by S-nitroso-N-acetylpenicillamine, a nitric oxide donorcompound. Int J InfectDis. 8: 223–226. [Crossref]

Khan, F. R., Syberg, K., Shashoua, Y., & Bury, N. R. (2015). Influence of polyethylene microplastic beads on the uptake and localization of silver in zebrafish (Danio rerio). Environmental pollution, 206, 73–79.

Kharazmi, A., Nielsen, H., Rechnitzer, C., & Bendtzen, K. (1989). Interleukin 6 primes human neutrophil and monocyteoxidativeburst response. Immunologyletters, 21(2), 177–184.

Klaassen, N., Spicer, V., & Krokhin, O. V. (2019). Universal retention standard for peptide separations using various modes of high-performance liquid chromatography. Journal of Chromatography A, 1588, 163–168.

Kolb, P., Ferreira, R. S., Irwin, J. J., & Shoichet, B. K. (2009). Docking and chemoinformatic screens for new ligands and targets. Current opinion in biotechnology, 20(4), 429–436.

Lamiable A, Thévenet P, Rey J, Vavrusa M, Derreumaux P, Tufféry P. PEP-FOLD3: previsão de estrutura de novo mais rápida para peptídeos lineares em solução e em complexo. NucleicAcids Res. 8 de julho de 2016; 44 (W1): W449–54.

Li, Y., Hu, Y., Yu, J., & Ma, T. (2020). Retrospectiveanalysisoflaboratorytesting in 54 patientswithsevere-orcritical-type 2019 novel coronavirus pneumonia. LaboratoryInvestigation, 1–7.

Liu, D., Thompson, J. R., Carducci, A., & Bi, X. (2020). Potential secondary transmission of SARS-CoV-2 via wastewater. Science of The Total Environment, 749, 142358.

Liu, H., Wu, J., Yao, J. Y., Wang, H., & Li, S. T. (2017). The role ofoxidative stress in decreasedacetylcholinesteraseactivity at the neuromuscular junctionofthediaphragmduring sepsis. Oxidative medicine and cellularlongevity, 2017.

Liu, W., Liu, Z., & Li, Y. C. (2020). COVID-19-related myocarditis and cholinergicanti-inflammatorypathways. HellenicJournalofCardiology.

Luna, O. F., Gomez, J., Cárdenas, C., Albericio, F., Marshall, S. H., & Guzmán, F. (2016). Deprotection reagents in Fmoc solid phase peptide synthesis: moving away from piperidine?. Molecules, 21(11), 1542.

Lusher, A. L., Mchugh, M., & Thompson, R. C. (2013). Occurrence of microplastics in the gastrointestinal tract of pelagic and demersal fish from the English Channel. Marine pollution bulletin, 67(1-2), 94–99.

Maharajan, K., Muthulakshmi, S., Nataraj, B., Ramesh, M., & Kadirvelu, K. (2018). Toxicity assessment of pyriproxyfen in vertebrate model zebrafish embryos (Danio rerio): a multi biomarker study. Aquatic Toxicology, 196, 132–145.

Mazloom, R. (2020). FeasibilityofTherapeuticEffectsoftheCholinergic Anti-InflammatoryPathwayon COVID-19 Symptoms. Journal of NeuroimmunePharmacology, 1–2.

Metzger, J. W., Kempter, C., Wiesmuller, K. H., & Jung, G. (1994). Electrospray mass spectrometry and tandem mass spectrometry of synthetic multicomponent peptide mixtures: determination of composition and purity. Analytical biochemistry, 219(2), 261–277.

Meyerowitz, E. A., Richterman, A., Gandhi, R. T., & Sax, P. E. (2020). Transmission of SARS-CoV-2: a review of viral, host, and environmental factors. Annals of internal medicine.

Milatovic, D., Gupta, R. C., & Aschner, M. (2006). Anticholinesterasetoxicity and oxidative stress. TheScientificWorldJournal, 6.

Miranda NEO, Maciel NM, Ribeiro MSL, Colli GR, Haddad FB, Collevatti RG. Diversification of the widespread neotropical frog Physalaemus cuvieri in response to Neogene-Quaternary geological events and climate dynamics. Molecular Phylogeneticsand Evolution, 132: 67–80, 2019.

Montalvão, M. F., Guimarães, A. T. B., de Lima Rodrigues, A. S., & Malafaia, G. (2021). Carbon nanofibers are bioaccumulated in Aphyllawilliamsoni (Odonata) larvae and cause REDOX imbalance and changes of acetylcholinesterase activity. Science of The Total Environment, 143991.

Nishikawa, M., Hashida, M., & Takakura, Y. (2009). Catalasedeliveryforinhibiting ROS-mediatedtissueinjury and tumor metastasis. Advanceddrugdeliveryreviews, 61(4), 319–326.

Ohkawa, H., Ohishi, N., & Yagi, K. (1979). Assay for lipid peroxides in animal tissues by thiobarbituric acid reaction. Analytical biochemistry, 95(2), 351–358.

Osman, A. H. (2020). COVID-19: Targetingthecytokinestormviacholinergicanti-inflammatory (Pyridostigmine). Int. J. Clin. Virol, 4, 041–046.

Osman, A. H. (2020). COVID-19: Targeting the cytokine storm via cholinergic anti-inflammatory (Pyridostigmine). Int. J. Clin. Virol, 4, 041–046.

Pais, F. S. M., de Cássia Ruy, P., Oliveira, G., & Coimbra, R. S. (2014). Assessing the efficiency of multiple sequence alignment programs. Algorithms for molecular biology, 9(1), 4.

Pala, A. (2019). Theeffectof a glyphosate-basedherbicideonacetylcholinesterase (AChE) activity, oxidative stress, and antioxidantstatus in freshwateramphipod: Gammaruspulex (Crustacean). EnvironmentalScience and PollutionResearch, 26(36), 36869–36877.

Pandey, D., Verma, S., Verma, P., Mahanty, B., Dutta, K., Daverey, A., & Arunachalam, K. (2020). SARS-CoV-2 in wastewater: Challenges for developing countries. International journal of hygiene and environmental health, 113634.

Pechmann, J. H., Scott, D. E., Semlitsch, R. D., Caldwell, J. P., Vitt, L. J., & Gibbons, J. W. (1991). Declining amphibian populations: the problem of separating human impacts from natural fluctuations. Science, 253(5022), 892–895.

Pettersen, EF, Goddard, TD, Huang, CC, et al. UCSF ChimeraX: Structure visualization for researchers, educators, and developers. Protein Science. 2021; 30: 70–82. https://doi.org/10.1002/pro.3943

Polo, D., Quintela-Baluja, M., Corbishley, A., Jones, D. L., Singer, A. C., Graham, D. W., & Romalde, J. L. (2020). Making waves: Wastewater-based epidemiology for SARS-CoV-2–Developing robust approaches for surveillance and prediction is harder than it looks. Water Research.

Polonikov, A. (2020). EndogenousDeficiencyofGlutathione as theMostLikely Cause ofSeriousManifestations and Death in COVID-19 Patients. ACS InfectiousDiseases.

Pupin, N. C., Gasparini, J. L., Bastos, R. P., Haddad, C. F., & Prado, C. (2010). Reproductive biology of an endemic Physalaemus of the Brazilian Atlantic forest, and the trade-off between clutch and egg size in terrestrial breeders of the P. signifer group. The Herpetological Journal, 20(3), 147–156.

Qi, X., Ke, B., Feng, Q., Yang, D., Lian, Q., Li, Z., … & Liao, G. (2020). Construction and immunogenic studies of a mFc fusion receptor binding domain (RBD) of spike protein as a subunit vaccine against SARS-CoV-2 infection. Chemical Communications, 56(61), 8683–8686.

Raibaut, L., El Mahdi, O., & Melnyk, O. (2014). Solid phase protein chemical synthesis. In Protein Ligation and Total Synthesis II (pp. 103–154). Springer, Cham.

Ranvestel, A. W., Lips, K. R., Pringle, C. M., Whiles, M. R., & Bixby, R. J. (2004). Neotropical tadpoles influence stream benthos: evidence for the ecological consequences of decline in amphibian populations. Freshwater Biology, 49(3), 274–285.

Ravichandran, S., Coyle, E. M., Klenow, L., Tang, J., Grubbs, G., Liu, S., … & Khurana, S. (2020). Antibody signature induced by SARS-CoV-2 spike protein immunogens in rabbits. Science Translational Medicine.

Ro, J. H., Liu, C. C., & Lin, M. C. (2020). Resveratrol Mitigates Cerebral Ischemic Injury by Altering Levels of Trace Elements, Toxic Metal, Lipid Peroxidation, and Antioxidant Activity. Biological Trace Element Research, 1–10.

Rutkoski, C. F., Macagnan, N., Folador, A., Skovronski, V. J., do Amaral, A. M., Leitemperger, J. W., … & Hartmann, M. T. (2020). Cypermethrin-and fipronil-based insecticides cause biochemical changes in Physalaemusgracilis tadpoles. Environmental Science and Pollution Research, 1–11.

Sachett, A., Bevilaqua, F., Chitolina, R., Garbinato, C., Gasparetto, H., Dal Magro, J., … & Siebel, A. M. (2018). Ractopamine hydrochloride induces behavioral alterations and oxidative status imbalance in zebrafish. Journal of Toxicology and Environmental Health, Part A, 81(7), 194–201.

Samrat, S. K., Tharappel, A. M., Li, Z., & Li, H. (2020). Prospect of SARS-CoV-2 spike protein: Potential role in vaccine and therapeutic development. Virus research, 198141.

Sangkham, S. (2020). Face mask and medical waste disposal during the novel COVID-19 pandemic in Asia. Case Studies in Chemical and Environmental Engineering, 2, 100052.

Santiago, I., Moreno-Munoz, A., Quintero-Jiménez, P., Garcia-Torres, F., & Gonzalez-Redondo, M. J. (2020). Electricity demand during pandemic times: The case of the COVID-19 in Spain. Energy policy, 148, 111964.

Sharma, H. B., Vanapalli, K. R., Cheela, V. S., Ranjan, V. P., Jaglan, A. K., Dubey, B., … & Bhattacharya, J. (2020). Challenges, opportunities, and innovations for effective solid waste management during and post COVID-19 pandemic. Resources, conservation and recycling, 162, 105052.

Shutler, J., Zaraska, K., Holding, T. M., Machnik, M., Uppuluri, K., Ashton, I., … & Dahiya, R. (2020). Risk of SARS-CoV-2 infection from contaminated water systems. MedRxiv.

Silva, F. F. D., Silva, J. M. D., Silva, T. D. J. D., Tenorio, B. M., Tenorio, F. D. C. A. M., Santos, E. L., … & Soares, E. C. (2020). Evaluation of Nile tilapia (Oreochromis niloticus) fingerlings exposed to the pesticide pyriproxyfen. Latin american journal of aquatic research, 48(5), 826–835.

Suhail, S., Zajac, J., Fossum, C., Lowater, H., McCracken, C., Severson, N., … & Bhattacharyya, S. (2020). Role ofOxidative Stress on SARS-CoV (SARS) and SARS-CoV-2 (COVID-19) Infection: A Review. Theproteinjournal, 1–13.

Tougu, V. (2001). Acetylcholinesterase: mechanism of catalysis and inhibition. Current Medicinal Chemistry-Central Nervous System Agents, 1(2), 155–170.

Tracey KJ. Physiology and immunologyofthecholinergicanti-inflammatorypathway. J Clin Invest. 2007; 117: 289–296.

Tran, H. N., Le, G. T., Nguyen, D. T., Juang, R. S., Rinklebe, J., Bhatnagar, A., … & Chao, H. P. (2020). SARS-CoV-2 coronavirus in water and wastewater: A critical review about presence and concern. Environmental Research, 110265.

Trott, O; Olson, A.J. AutoDock Vina: improving the speed and accuracy of docking with a new scoring function, efficient optimization, and multithreading. J Comput Chem. 2010;31(2):455–461. doi:10.1002/jcc.21334

Tsujimoto, M., Yokota, S., Vilček, J., & Weissmann, G. (1986). Tumor necrosis factor provokessuperoxideaniongenerationfromneutrophils. Biochemical and biophysicalresearchcommunications, 137(3), 1094–1100.

Urban, R. C., & Nakada, L. Y. K. (2021). COVID-19 pandemic: Solid waste and environmental impacts in Brazil. Science of the Total Environment, 755, 142471.

Valavanidis, A., Vlahogianni, T., Dassenakis, M., & Scoullos, M. (2006). Molecular biomarkers of oxidative stress in aquatic organisms in relation to toxic environmental pollutants. Ecotoxicology and environmental safety, 64(2), 178–189.

Wang, X. W., Li, J. S., Jin, M., Zhen, B., Kong, Q. X., Song, N., … & Si, B. Y. (2005). Study on the resistance of severe acute respiratory syndrome-associated coronavirus. Journal of virological methods, 126(1-2), 171–177.

Waterhouse, A., Bertoni, M., Bienert, S., Studer, G., Tauriello, G., Gumienny, R., Heer, F.T., de Beer, T.A.P., Rempfer, C., Bordoli, L., Lepore, R., Schwede, T. SWISS-MODEL: homology modelling of protein structures and complexes. Nucleic Acids Res. 46(W1), W296–W303 (2018).

Wu, F., Xiao, A., Zhang, J., Moniz, K., Endo, N., Armas, F., … & Duvallet, C. (2020). SARS-CoV-2 titers in wastewater foreshadow dynamics and clinical presentation of new COVID-19 cases. Medrxiv.

Wrubleswski, J., Reichert, F. W., Galon, L., Hartmann, P. A., & Hartmann, M. T. (2018). Acute and chronic toxicity of pesticides on tadpoles of Physalaemus cuvieri (Anura, Leptodactylidae). Ecotoxicology, 27(3), 360–368.

Xiao, F., Sun, J., Xu, Y., Li, F., Huang, X., Li, H., … & Zhao, J. (2020). Infectious SARS-CoV-2 in feces of patient with severe COVID-19. Emerging infectious diseases, 26(8), 1920.

Yang, J., Wang, W., Chen, Z., Lu, S., Yang, F., Bi, Z., … & Hong, W. (2020). A vaccine targeting the RBD of the S protein of SARS-CoV-2 induces protective immunity. Nature, 586(7830), 572–577.

Yang, L., Yu, X., Wu, X., Wang, J., Yan, X., Jiang, S., & Chen, Z. (2020). Emergency response to the explosive growth of health care wastes during COVID-19 pandemic in Wuhan, China. Resources, Conservation and Recycling, 164, 105074.

Zand, A. D., & Heir, A. V. (2020). Emerging challenges in urban waste management in Tehran, Iran during the COVID-19 pandemic. Resources, Conservation, and Recycling, 162, 105051.

